# A uniquely stable trimeric model of SARS-CoV-2 spike transmembrane domain

**DOI:** 10.1101/2022.06.05.494856

**Authors:** E.T. Aliper, N.A. Krylov, D.E. Nolde, A.A. Polyansky, R.G. Efremov

## Abstract

The spike (S) protein of SARS-CoV-2 effectuates membrane fusion and virus entry into target cells. Its transmembrane domain (TMD) represents a homotrimer of α-helices anchoring the spike in the viral envelope. Although S-protein models available to date include the TMD, its precise configuration was given brief consideration. Understanding viral fusion entails realistic TMD models, while no reliable approaches towards predicting the 3D structure of transmembrane (TM) trimers exist. Here, we propose a comprehensive computational framework to model the spike TMD (S-TMD) based solely on its primary structure. First, we performed amino acid sequence pattern matching and compared molecular hydrophobicity potential (MHP) distribution on the helix surface against TM homotrimers with known 3D structures and thus selected the TMD of the tumour necrosis factor receptor 1 (TNFR-1) for subsequent template-based modelling. We then iteratively built an all-atom homotrimer model of S-TMD based on “dynamic MHP portraits” and residue variability motifs. In this model each helix possessed two overlapping interfaces interacting with either of the remaining helices, which include conservative residues I1216, F1220, I1227, M1229, and M1233. Finally, the stability of this and several alternative models (including a recent NMR structure) and a set of mutant forms was tested in all-atom molecular dynamics (MD) simulations in a POPC bilayer mimicking the viral envelope membrane. Unlike other configurations, our model trimer remained extraordinarily tightly packed over a microsecond-range MD and retained its stability when palmitoylated in accordance with experimental data. Palmitoylation had no significant impact on the TMD conformation nor the way in which the lipid bilayer was perturbed in the presence of the trimer. Overall, the resulting model of S-TMD conforms to known basic principles of TM helix packing and will be further used to explore the complex machinery of membrane fusion from a broader perspective beyond the TMD.

## INTRODUCTION

The spike (S) protein, crucial to the infectivity of SARS-CoV-2 and other coronaviruses, is a class I viral fusion protein, alongside a number of proteins that have long been under scrutiny including HIV’s gp-41 and haemagglutinin from influenza virus. Like other fusogens of this kind, the S-protein is trimeric and consists of a voluminous ectodomain exposed on the virion surface, an α-helical transmembrane domain (TMD) and a small endodomain (Cai et al., 2020). Apart from its well-documented role in receptor recognition, the spike, or, more specifically, its S2 subunit, is a factor effectuating membrane fusion, bringing together the viral and target membranes, facilitating the release of the viral genome into the target cell. To accomplish this function, fusogenic proteins employ their regions tailored to interact with the membranes of both the viral and host cell - TMD(s) and one or several membrane-active fragments such as the fusion peptides (Xia et al., 2020; Basso et al., 2021). Therefore, in order to understand all stages of viral fusion, one would require exhaustive knowledge of the TMD’s structural subtleties, as they directly contribute to spike protein refolding and the merging of the membranes.

Little experimental information on the SARS-CoV-2 TMD structure has been obtained so far. Recently, a trimerization study has been conducted by Fu and Chou (Fu and Chou, 2021) for a peptide corresponding to SARS-CoV-2’s spike residues 1209-1237, in which Met and Cys residues were substituted for Leu and Ser, respectively, amounting to a total of four point mutations compared to the wild type protein. In dimyristoylphosphatidylcholine / 1,2-Dihexanoyl-sn-Glycero-3-Phosphocholine (DMPC/DH_6_PC) bicelles this peptide assumed a trimeric structure with a Leu/Ile-zipper-like interface. The portion of the peptide whereof the structure was resolved and which spanned the bicelle membrane was mapped as residues 1218 to 1234, corresponding to a rather short TMD fragment 16 residues long (PDB ID 7LC8). Immediately downstream of its ectodomain the spike contains a small region rich in aromatic residues (1212-WPWYIW-1217), followed by a hydrophobic region (1218-LGFIAGLIAIVMVMTIML-1234). It has previously been shown in an NMR-based study (Mahajan and Bhattacharjya, 2015) that a peptide corresponding to the C-terminal portion of the stem region followed by the aromatic cluster of the spike in SARS-CoV (residues 1193 to 1202), whereof the spike protein S2 subunit is highly homologous to that of SARS-CoV-2, has the propensity to assume a helix-loop-helix conformation in dodecylphosphocholine micelles, the loop roughly coinciding with the area around I1210 and K1211 (SARS-CoV-2 numbering). Four clusters of at least two cysteines can be found downstream of the TMD, the first one located immediately after it and consisting of Cys 1235 and Cys1236. Both of these residues have been shown to be palmitoylated, and this modification is believed to be crucial to fusion, as replacement of these residues with alanine resulted in a considerable loss of infectivity, in both SARS-CoV-2 (Mesquita et al., 2021) and SARS-CoV (Petit et al., 2007).

Despite experimental structural data on the organisation of the spike TMD being scarce, several attempts have been made to model it *in silico*. A number of full-length models of SARS-CoV-2’s spike protein have been built, within which the TMD was reconstructed using template-based modelling. However, it was on no occasion the primary focus of modelling efforts, and the finer, more intricate aspects of its structure might have been overlooked. In one of these models, created by Casalino et al. (2020), a water-soluble coiled-coil from a bacterial protein was used as the template, although transmembrane (TM) α-helices are known to possess a primary structure only peculiar to them to guarantee optimum packing tailored to the lipid environment (Rees et al., 1989; Lemmon and Engelman, 1992). This model only includes palmitoyl modifications at Cys1240 and Cys1241, known as the second cysteine cluster. Shortly afterwards, another model became available (Woo et al., 2020), within which the TMD had been built via homology modelling using NMR data obtained for the TMD from HIV’s gp41, another class I fusion protein and hence the spike’s close functional counterpart. The same structure was used for the template-based modelling of S-TMD in a recent *in silico* study of allosteric effects in the spike (Tan et al., 2022). However, it was not examined in detail to what extent the physico-chemical properties of the TM helix were similar in the two fusion proteins. Nor has it been revealed whether the proposed model is in agreement with the known principles of helical TMD organisation, namely its structural aspect as well as the distribution of hydrophobic/hydrophilic and conservative/variable residues between the lumen of the trimer and lipid-exposed surface. The models available for downloading do not include palmitoyl modifications, however, the authors used CHARMM-GUI (Jo et al., 2008) to add palmitoyl tails to Cys1236 and Cys1241 to perform MD simulations of full-length spike protein binding to its receptor and antibodies. Other models have been derived relying on various automated tools such as Rosetta and I-TASSER as opposed to template-based modelling (ZhangLab, 2020; Izvorski, 2020; Nishima and Kulik, 2021), however, it is not possible to evaluate their overall quality at this time.

Like in the case of other integral membrane proteins, the main problem in deciphering the spatial structure of TMDs of viral fusogens is that they have to be stabilised in a native-like membrane environment, while much more voluminous domains of the spike need to stay in water. Also, in the immediate vicinity of the TMD there are several very conformationally flexible membrane-active regions, which can hamper efforts to obtain a high-resolution structure using X-ray crystallography or cryo-EM techniques. From the computational point of view, the problem lies in the absence of reliable instruments for the prediction of TM homotrimer 3D structure. In the present study, we explore the possibility of solving this task *in silico* for the TMD of SARS-CoV-2’s spike protein (S-TMD) solely on the basis of the amino acid sequence. Our main objective was to construct such a model taking into account all the known principles of TM helix packing identified so far. To accomplish this, we designed an original computational framework that is described below.

## RESULTS

### Modelling framework

The design of the present study was as follows:

1. At the first stage, the **boundaries of S-TMD were identified** via **sequence analysis** and further ascertained via **Monte Carlo (MC) simulations** of a TM monomer in an implicit membrane. Eventually, the fragment chosen to model S-TMD consisted of residues 1212 to 1234.
2. At the next stage, **an optimal structural template was selected** among homotrimers of TM helices with known structure. The template had to conform to the following two criteria: (1) the presence of patterns of residues with physicochemical properties in the TM sequence sufficiently similar to those in the fragment of interest in spike; (2) a good correspondence of the surface geometric and MHP properties between the two TM segments (in the template and spike). In the absence of sequence homology with available potential templates, the main assumption was that TM helices in S-TMD would pack in a similar way as in a template whereof the monomers possess similar sequence/surface motifs. To this end we scrutinised the TM sequence of the spike and compared it to all TM homotrimers in the PDB database in search of common patterns of charged, polar, hydrophobic, small and proline residues. For the candidates thus identified, we compared spatial distributions of hydrophobic/hydrophilic properties on their solvent-accessible surface and selected the optimal one(s).
3. To enhance the reliability of S-TMD modelling, we used an independent approach. On the basis of MHP complementarity for TM helices, we predicted dimerization interfaces for the TM segment in order to identify surfaces likely to be on the helix-helix interfaces and compared them against those in the template(s) to pick the optimum one, with helix/helix contact areas as similar to those predicted for S-TMD as possible.
4. One of the candidate templates, the TMD of TNFR-1, was chosen to **build a model of S-TMD** via homology modelling. To **test the stability** and viability of this model, **MD simulations** of this trimer were performed in a model POPC bilayer. POPC was chosen, as it mimics, to a sufficient extent, the thickness and composition of the membrane of the endoplasmic reticulum Golgi intermediate compartment, where SARS-CoV-2 virions acquire their envelopes. Additionally, MD simulations of the original template were performed to evaluate its behaviour in a POPC bilayer. Based upon MD data a new, better packed model trimer (*S_OPT*) was built via iterative adjustment of the ‘dynamic MHP portraits’ of the interacting helices.
5. The **stability** of model trimer *S_OPT* and several alternative ones was then **evaluated** via MD simulations (see Tables 1 and 2).
6. Finally, **palmitoyl tails** were **added** at C1235 and C1236 in the *S_OPT* model in order to explore **how such modifications could affect the stability** of S-TMD and the way it was accommodated by the membrane during MD simulations.

**Table 1.**
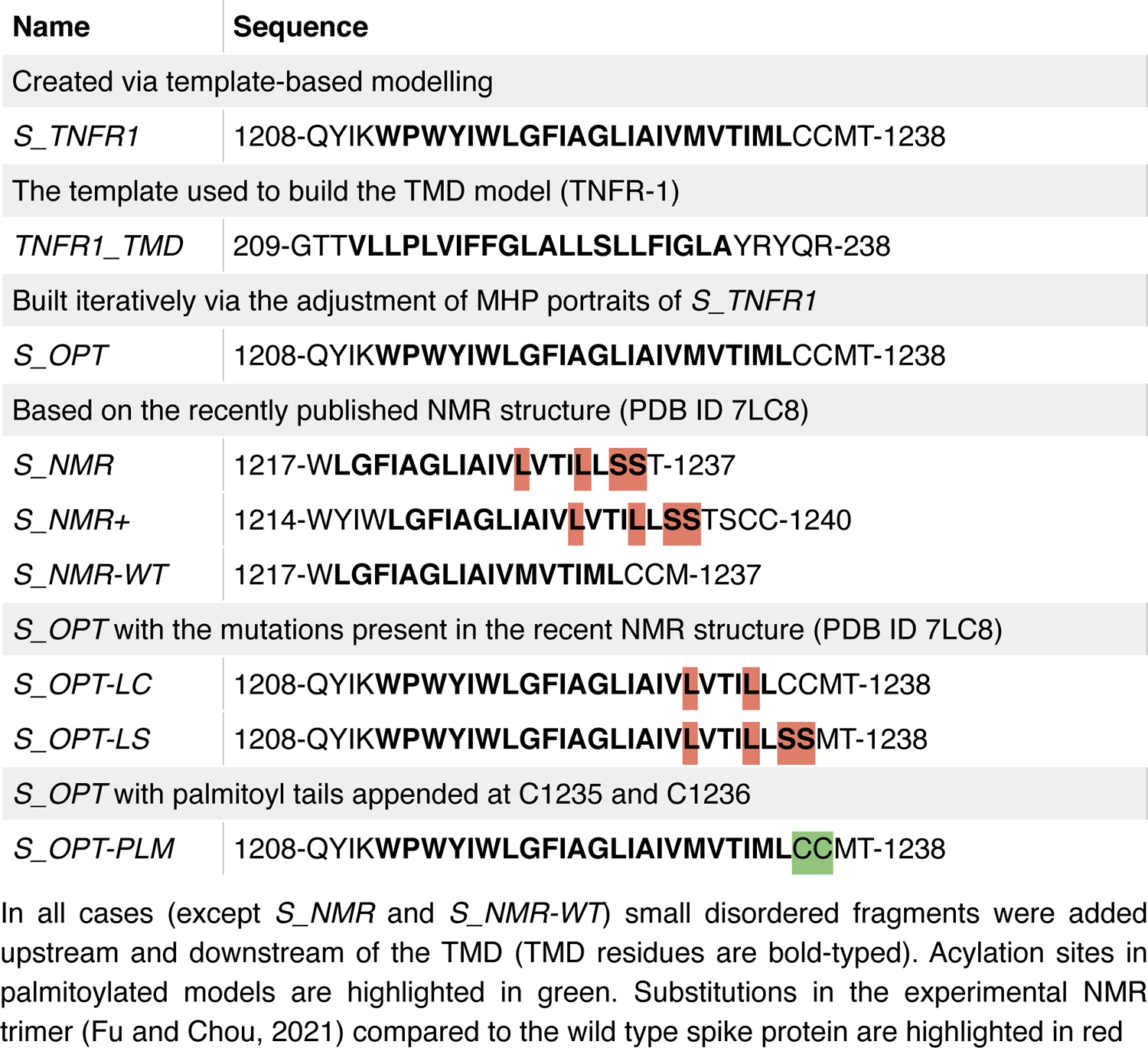
Model trimers tested in the present study.

**Table 2.** MD trajectories calculated in the present study.

| System composition | MD length<br>(ns) | Number of<br>replicas |
| --- | --- | --- |
| *A POPC bilayer* |  |  |
| POPC<sub>450</sub> / Water<sub>26276</sub> | 1000 | 1 |
| *TNFR1_TMD in a lipid bilayer* |  |  |
| Protein / POPC<sub>475</sub> / Water<sub>42057</sub> / Cl<sup>-</sup><sub>6</sub> | 500 | 1 |
| *S_TNFR1 in a lipid bilayer* |  |  |
| Protein / POPC<sub>483</sub> / Water<sub>42060</sub> / Cl<sup>-</sup><sub>3</sub> | 500-1000 | 4 |
| *S_OPT in a lipid bilayer* |  |  |
| Protein / POPC<sub>485</sub> / Water<sub>42098</sub> / Cl<sup>-</sup><sub>3</sub> | 1000 | 2 |
| *structures based on the experimentally observed trimer (PDB ID 7LC8) in a lipid bilayer* |  |  |
| *S_NMR* / POPC<sub>488</sub> / Water<sub>42196</sub> | 500 | 1 |
| *S_NMR+* / POPC<sub>485</sub> / Water<sub>42148</sub> | 500 | 1 |
| *S_NMR-WT* / POPC<sub>485</sub> / Water<sub>42148</sub> | 500 | 1 |
| *S_OPT with the mutations present in the experimentally observed trimer (PDB ID 7LC8) in a lipid bilayer* |  |  |
| *S_OPT-LC* / POPC<sub>483</sub> / Water<sub>42092</sub> / Cl<sup>-</sup><sub>3</sub> | 1000 | 1 |
| *S_OPT-LS* / POPC<sub>484</sub> / Water<sub>42117</sub> / Cl<sup>-</sup><sub>3</sub> | 1000 | 1 |
| *S_OPT palmitoylated at C1235 and C1236* |  |  |
| Protein / POPC<sub>480</sub> / Water<sub>42084</sub> / Cl<sup>-</sup><sub>3</sub> (run1) | 1000 | 1 |
| Protein / POPC<sub>474</sub> / Water<sub>42120</sub> / Cl<sup>-</sup><sub>3</sub> (run2) | 1000 | 1 |

The number of molecules/ions of each type in the system, if above 1, is presented in subscript. See Table 1 for more details on the peptides studied.

### S-TMD boundaries identified via sequence analysis and Monte Carlo simulations

Different TMD boundaries were predicted with recourse to TMSEG (TMD residues: 1214-1238) and TMPRED (1216-1235) or proposed on the basis of data from UniProt entry P0DTC2 (1214-1234), as well as in the work by Xia et al. (2020) (1213-1237) and Cai et al. (1212-1234) (2020). Subsequent adjustment of S-TMD’s boundaries was done using MC simulations of an S-TMD monomer (residues 1208 to 1239) in an implicit membrane. It should be noted that this segment encompasses all the regions mentioned above.

Analysis of the conformers accumulated in the course of MC search invites the following conclusions: 1) Throughout the entire simulation the peptide preserves its initial α-helical structure well in the portion formed by residues from I1216 to L1234; 2) Being initially placed in the water phase, it partially penetrates into the hydrophobic layer (ǀZǀ < 15 Å) as early as upon preliminary minimization, before the actual MC steps were performed. 3) Subsequent conformational search in the dihedral angles space was accompanied by a decrease of the total energy (*Etot.*) of the system, during which the peptide quickly assumed the TM orientation. 4) The ensemble of the accumulated low-energy MC-states (within 5 kcal/mol from the minimal *Etot.*) only includes the TM segment in the α-helical conformation (residues I1216 to L1234), which spans the hydrophobic layer mimicking the “membrane” (Fig. 1). In these states, the angle between the helical axis and the membrane normal (Z-axis) is 25° ± 2°. The peptide is kinked at residue Y1215, while its N-terminal part (residues 1208 to 1215) is disordered. In these states residues I1210 and W1212 to Y1215 are buried in the membrane (Fig. 1).

**Figure 1.**
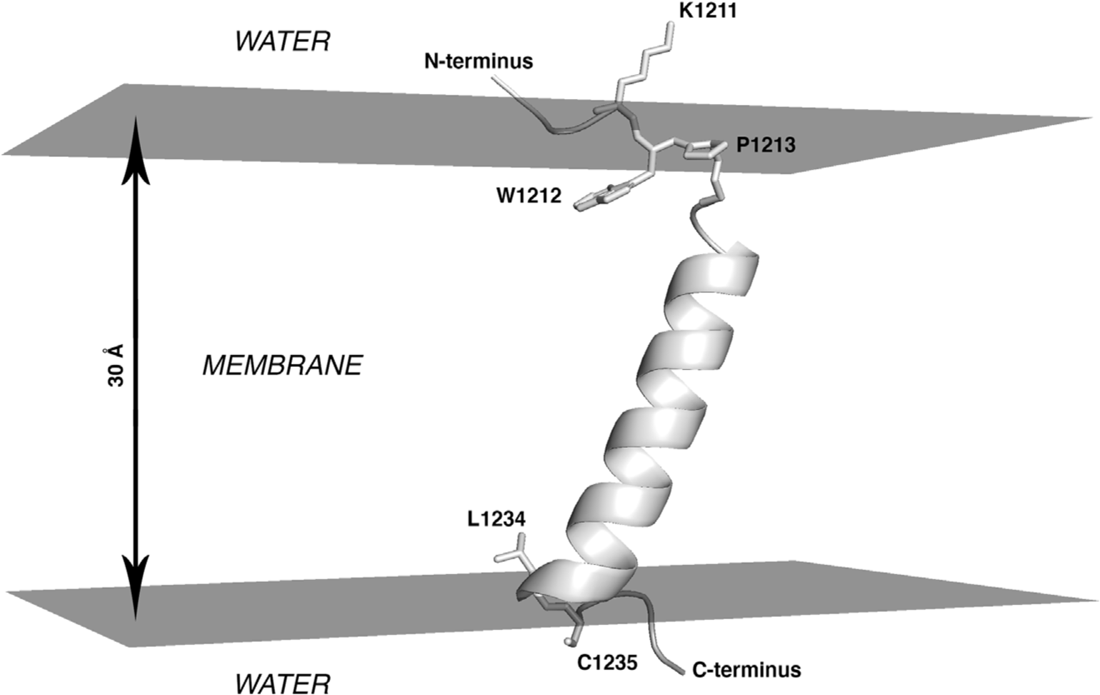
Lowest-energy state for monomeric S-TMD derived via Monte Carlo simulations. Spike fragment 1208-1239 in an implicit membrane is shown. The system is oriented in such a way that the top and bottom surfaces of the hydrophobic slab (here represented in grey) are perpendicular to the Z axis and correspond to XY planes with Z-coordinates of +15Å and −15Å. The protein is shown in cartoon representation with selected residues in stick representation.

This result indicates that in the most energetically favourable MC-states the entire TMS peptide is embedded into the hydrophobic medium and does not expose the N-terminal fragment from W1212 to Y1215 at the lipid/water interface. A different orientation has been proposed in other studies (Fu and Chou, 2021), in which the region between Y1209 and W1217 was considered to be a juxtamembrane domain, while the TM part only included residues L1218 to L1234. MC data therefore additionally justify the selection of the entire fragment from W1212 to L1234 to build a model of S-TMD trimer. This choice is also supported by the fact that K1211 immediately upstream thereof is a charged residue highly unlikely to be part of the TMD, and residues 1235 and 1236 are palmitoylated cysteines, likely to be membrane-proximal, but not belonging to the TMD *per se*.

### TMD of tumour necrosis factor receptor-1 as an optimal template for S-TMD modelling

Analysis of sequence patterns formed by polar, hydrophobic, small and proline residues in a set of homotrimeric TMDs with known 3D structure (Table S1) revealed two structures of interest. Selected as candidate templates for the modelling of S-TMD were lysosome-associated membrane protein 2A (LAMP-2A, PDB ID 2MOM (Rout et al., 2014)) and tumour necrosis factor receptor-1 (TNFR-1, PDB ID 7K7A (Zhao et al., 2020)). Corresponding sequence alignment is presented in Fig. 2A. In spike, TNFR-1 and LAMP-2 one can notice a hydrophobic N-terminal residue followed by a proline and several sporadically located small residues. The polar serine in TNFR-1 was known to not be located on the helix/helix interface, which would therefore be constituted by small and hydrophobic side chains. Close attention was also paid to the TMD of HIV’s gp41 protein (Dev et al., 2016), a class I viral fusion protein and thus spike’s functional counterpart (PDB ID 5JYN). However, its sequence features two charged arginine residues (Fig. 2A), resulting in a pattern different from that in S-TMD, which led us to focus on other candidate templates.

**Figure 2.**
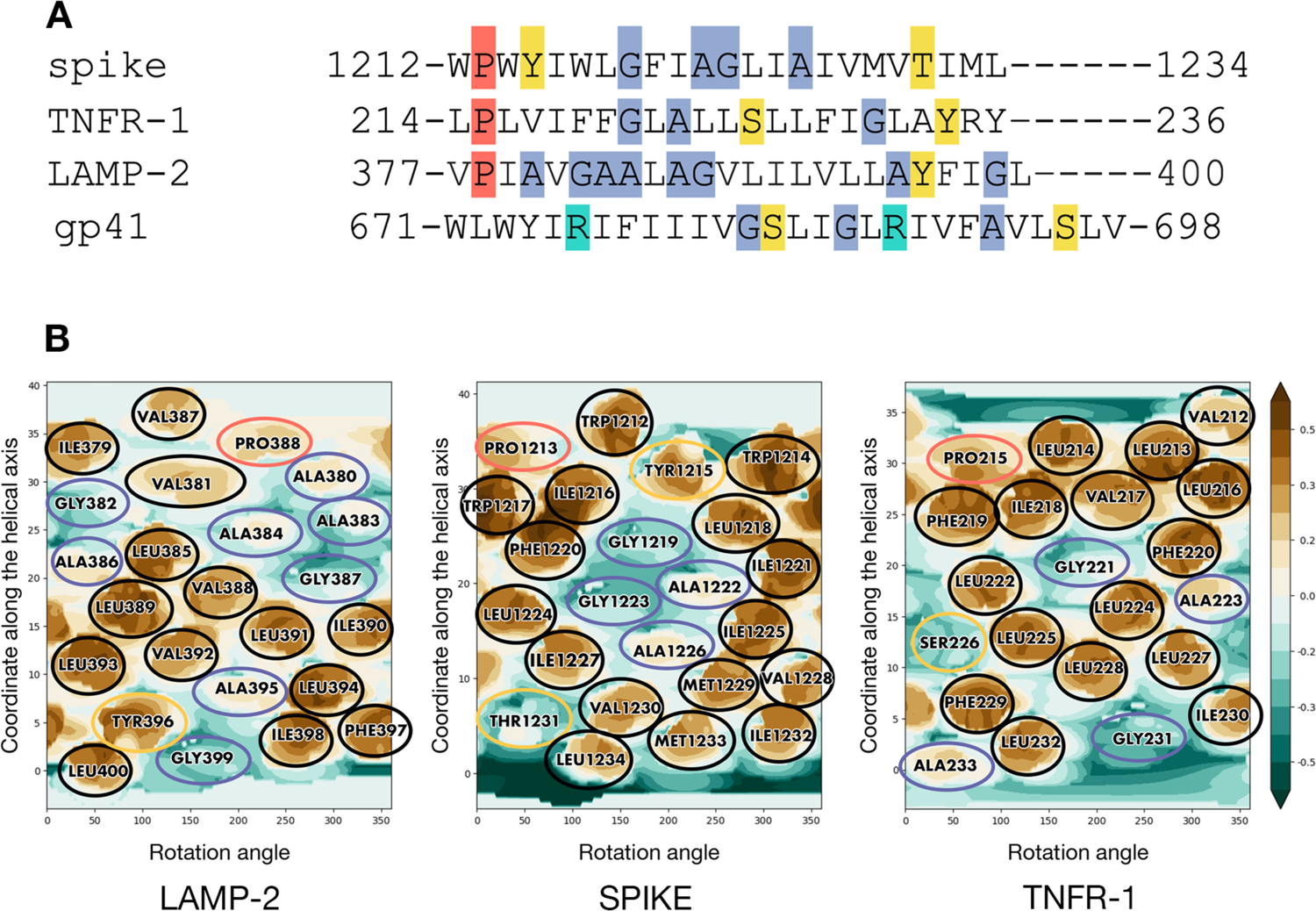
Residue patterns in S-TMD and candidate templates. (A) Similarity between patterns in the sequences of spike glycoprotein and TMDs with known 3D structure - LAMP-2 (2MOM), TNFR-1 (7K7A) and gp41 (5JYN). Hydrophobic residues are unhighlighted, while proline, small, charged and polar residues are highlighted in coral, blue, turquoise and yellow, respectively. (B) Molecular hydrophobicity potential (MHP) distribution maps for an ideal helix corresponding to S-protein residues 1212-1234 and TMD monomers of two candidate templates, LAMP-2 and TNFR-1. Cylindrical projection of the surface MHP distribution is used. Axis values correspond to the rotation angle around the helical axis and the coordinate along the latter (in Å), respectively. An MHP scale (in logP octanol-1/water units) is presented on the right. The maps are coloured in accordance with the MHP values (Efremov et al., 1992), from teal (hydrophilic areas) to brown (hydrophobic ones). Projections of proline, small, polar and hydrophobic residues are encircled in coral, blue, yellow and black, respectively.

2D MHP maps of monomeric TM helices for the two candidate templates (taken from the corresponding 3D models) and an ideal helix with a sequence corresponding to residues 1212-1234 in S-TMD are shown in Fig. 2B. It can be seen that the spike shares a common MHP pattern with TNFR-1: an islet of small residues (G1219, A1222, G1223, A1226 in spike and G221 in TNFR-1) surrounded by bulkier hydrophobic residues on all sides (Y1215, I1216, L1218, F1220, I1221, L1224, I1225, I1227, V1228, M1229 in spike and V217, I218, F219, F220, L222, L224, L225, L228 in TNFR-1). LAMP-2 has a similar small residue islet (A395 and G399), however, it lacks hydrophobic “lining” C-terminally, as it is immediately followed by the highly polar cytoplasmic domain, while in TNFR-1 this MHP motif is located “higher” up the helix, towards the middle of the TMD. In the light of these observations, we decided to use TNFR-1 as template in the modelling of S-TMD.

### Preliminary model of S-TMD homotrimer based on predicted helix-helix interfaces

On the basis of MHP complementarity between the surfaces of TM segments, three helix/ helix dimerization interfaces were identified for S-TMD and dubbed interface A, B and AB (Fig. S1). Amino acid positions in S-TMD were considered identical and (semi-)conservative if they were such among diverse members of genus *Betacoronavirus*. Apart from this, helix-helix interfaces for the TMD of the template TNFR-1 were evaluated using its spatial structure. It turned out to be organised as follows: each helix has two partially overlapping interfaces, interface 1 and interface 2, interacting with either of the two remaining helices. In every helix, interface 1 interacts with interface 2 and vice versa (Fig. 3C).

**Figure 3.**
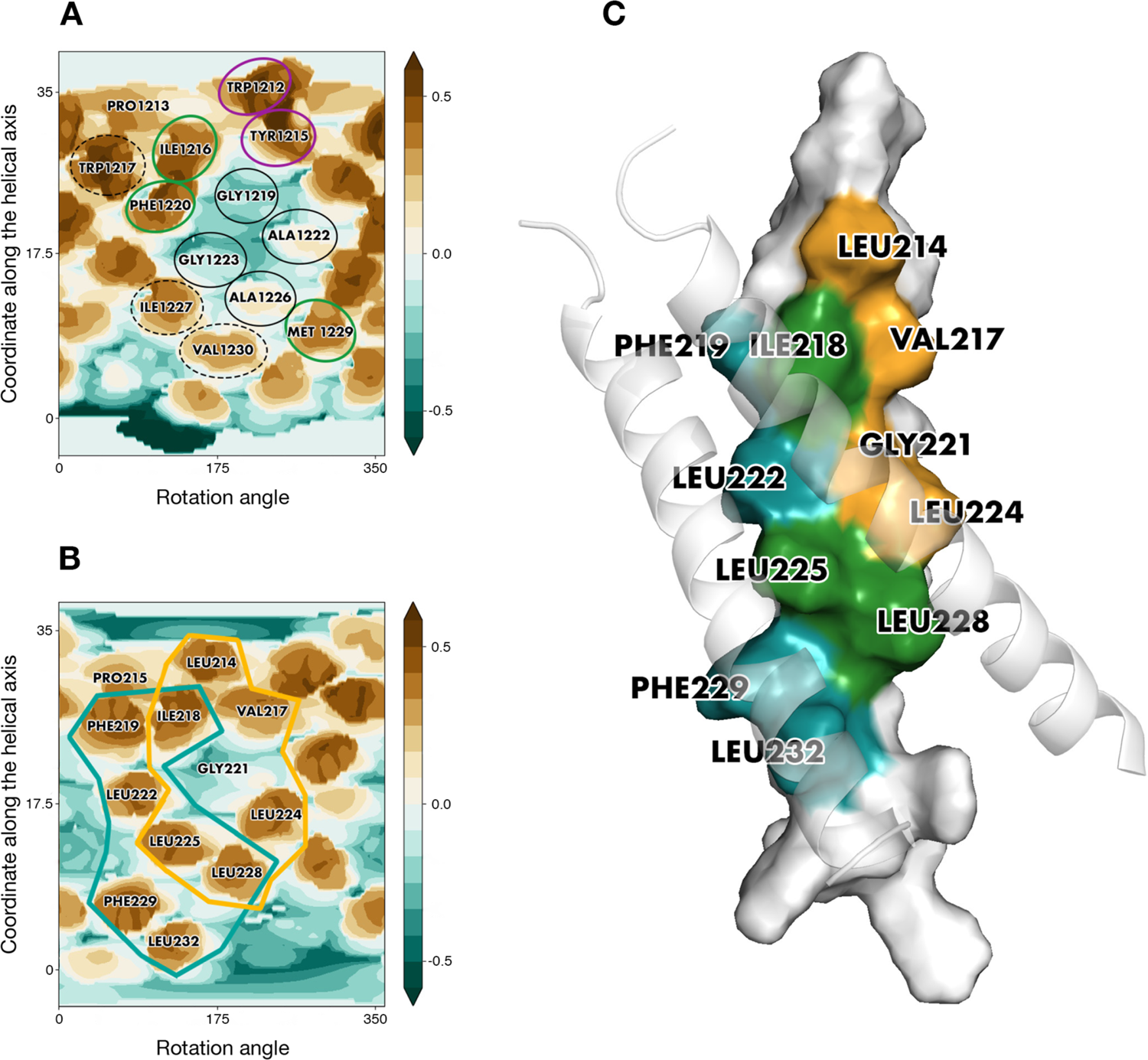
Similarity between S-TMD and a candidate template. (A) Interface B, one of dimerization solutions predicted for S-TMD. (B) Overlapping helix-helix interfaces for one of the chains in TNFR-1. TNFR-1’s interface 1 and interface 2 are enclosed in golden and teal lines, respectively. Identical positions are encircled in purple, conservative and semi-conservative residues are encircled in green, non-conservative residues present on the helix/helix interface are encircled in black, and residues in S-TMD that could be part of the helix-helix interface if the latter is similar to that of TNFR-1 are enclosed in dashed lines. For other details see legend to Fig. 2B. (C) A 3D model of the TMD of TNFR-1. Residues that are part of interface 1, part of interface 2 and part of both interfaces are coloured golden, teal and green, respectively. One of the helices is rendered in surface representation, while the other two helices are shown in cartoon representation and are semi-transparent.

Noteworthily, a pattern formed by W1212, Y1215, I1216, G1219, A1222, G1223 and A1226 in interface B of S-TMD was remarkably redolent of interface 1 identified for TNFR-1 (Fig. 3 A, B). Furthermore, if the two sequences were aligned for these two interfaces to match, proline residues in TNFR-1 and the ideal helix corresponding to spike’s residues 1212-1234 ended up in highly similar positions in relation to the helix/helix interface. Of equal interest was the fact that the ideal helix with the sequence of S-TMD had additional residues, W1217, F1220, I1227 and V1230, which, together with interface B added up to a pattern we had observed for the candidate template: two partially overlapping interfaces fit to accommodate two helices in a manner similar to that of TNFR-1, which was accordingly chosen for template-based modelling of the spike protein’s TMD.

### S_OPT: a tightly packed TMD model obtained via iterative refinement

Some differences were observed between TNFR-1 and our model of the spike protein’s TMD, *S_TNFR1*. Most notably, a hydrophobic patch on TNFR-1’s helix-helix interface constituted by one glycine (221) and three leucines (222, 224, 225) corresponded to two alanines (1222, 1226) and two glycines (1219, 1223) in our model. In line with this, the difference in solvent-accessible surface area, or ASA, (dASA) between *S_TNFR1* and three helices before assembly equalled ∼1500 Å^2^. In our template, TNFR-1, dASA was nearly twice as high, ∼2950 Å^2^, suggesting that a hollow space was present inside our model, a number of lumen-facing side chains located too far apart to interact and stabilise the trimer and not contributing to helix/helix interface formation (Fig. S2 D).

In the course of MD simulations of *S_TNFR1*, the number of protein-lipid (P/L) contacts grew dramatically, indicating that the helix/helix interface was at least partially exposed to POPC molecules and disrupted by them (Fig. S2 A). This is barely surprising, considering the imperfections of the trimer derived via direct template-based modelling. In some trajectories we observed the formation of a symmetrical dimer by two of the three helices of the model with helix/helix interfaces similar to the template’s predicted interface 2 and including I1227 and/or V1230, confirming that these residues are indeed fit to be located on helix/helix interfaces (data not shown). More interestingly yet, among the resulting MD-states we found a dimer formed by two of the three helices within the model trimer, whereof residues predicted to constitute interface 1 belonging to one helix interacted with residues predicted to be part of interface 2 belonging to the other helix (Fig.S2 E). This dimer formed as early as 9 ns and, in the microsecond range, proved to be stable and tightly packed, as is evidenced by RMSD (Fig. S2 B) and ASA (Fig. S2 C) dynamics over the course of the MD. dASA for this dimer is estimated at ∼880 Å^2^ (as calculated at 200 ns). At ∼830 ns, however, the dimer underwent a distinct change (Fig. S2 B) from asymmetrical to roughly symmetrical. After the M1229 in chain C moved away from the interface, so did its W1212, whilst its F1220 ended up facing chain A after slight rotation around the helical axis. Eventually roughly the same residues in both chains contributed to the helix/helix interface, albeit still undisrupted by lipids. This shift is evident from both the RMSD and ASA dynamics (Figs. S2 B and C). At the same moment in time (∼830 ns), the average number of pairs of atoms belonging to the side chains on the “initial” asymmetrical interface between helices A and С engaged in protein-protein (P/P) interactions also decreased from 23 to 21.

We thus obtained a structure that could potentially be used to build a *bona fide* trimer, organised in accordance with the same fundamental principles as the TMD of TNFR-1, but without noticeable empty spaces inside the helix bundle. To verify whether this was indeed the case, we manually “cloned” two trimers on the basis of the aforementioned asymmetrical dimer, aligning the “original” dimer and its copy in such a way as to obtain the missing third chain in the predicted position and later deleting the “extra” chain in the copy. The resulting two trimers were exceedingly similar to each other, and one, named *S_OPT*, was chosen for further examination.

Helix/helix interface maps in *S_OPT* were highly similar to those of TNFR-1, as though a post-packing event had occurred during simulation in a bilayer to perfect the structure and squeeze the helices closer together where the hollow space used to be. This space unstabilised by intermolecular interactions inside *S_TNFR1* had been partially located around the area (Fig. 4A) constituted by small residues (G1219, A1222, G1223, A1226) in lieu of a bulkier core of three leucines (222, 224, 225) and glycine 221 in TNFR-1. In addition to this, V1230 had also taken the place of leucine residues, failing to fill all of the volume fit to accommodate bulkier leucine chains. In *S_OPT* all of these residues were brought close together so that they ended up interacting (Fig. 4B). Furthermore, TNFR-1 had glycine residues at position 231, which in our model corresponded to M1229, more voluminous hydrophobic entities. Together with M1233 and L1234, corresponding to positions in TNFR-1 that are not even part of its transmembrane domain, R235 and Y1236, M1229 formed another tightly packed patch in *S_OPT*. This patch fits in well with the “two interfaces per helix, one to interact with either of the remaining two helices” principle, whereby the trimer gains additional turns of the helix contributing to the stabilisation of the interface. Much in the same vein, dASA for *S_OPT* equalled ∼2580 Å^2^, as opposed to ∼1500 Å^2^ in *S_TNFR1*, while the decrease of FV observed in the lumen of the trimer in *S_OPT* compared to the starting structure is estimated to be approximately 25-fold (Fig. 5). The empty space was thus eliminated, as the helices were positioned closer together and engaged in more favourable intermolecular P/P contacts, which stabilised the trimer. Accordingly, in the course of subsequent MD simulations (see the next section for details) *S_OPT* demonstrated much lower RMSD values as compared to *S_TNFR1*, amounting to 1.3±0.2 Å for the former and to 2.9±0.4 for the latter.

**Figure 4.**
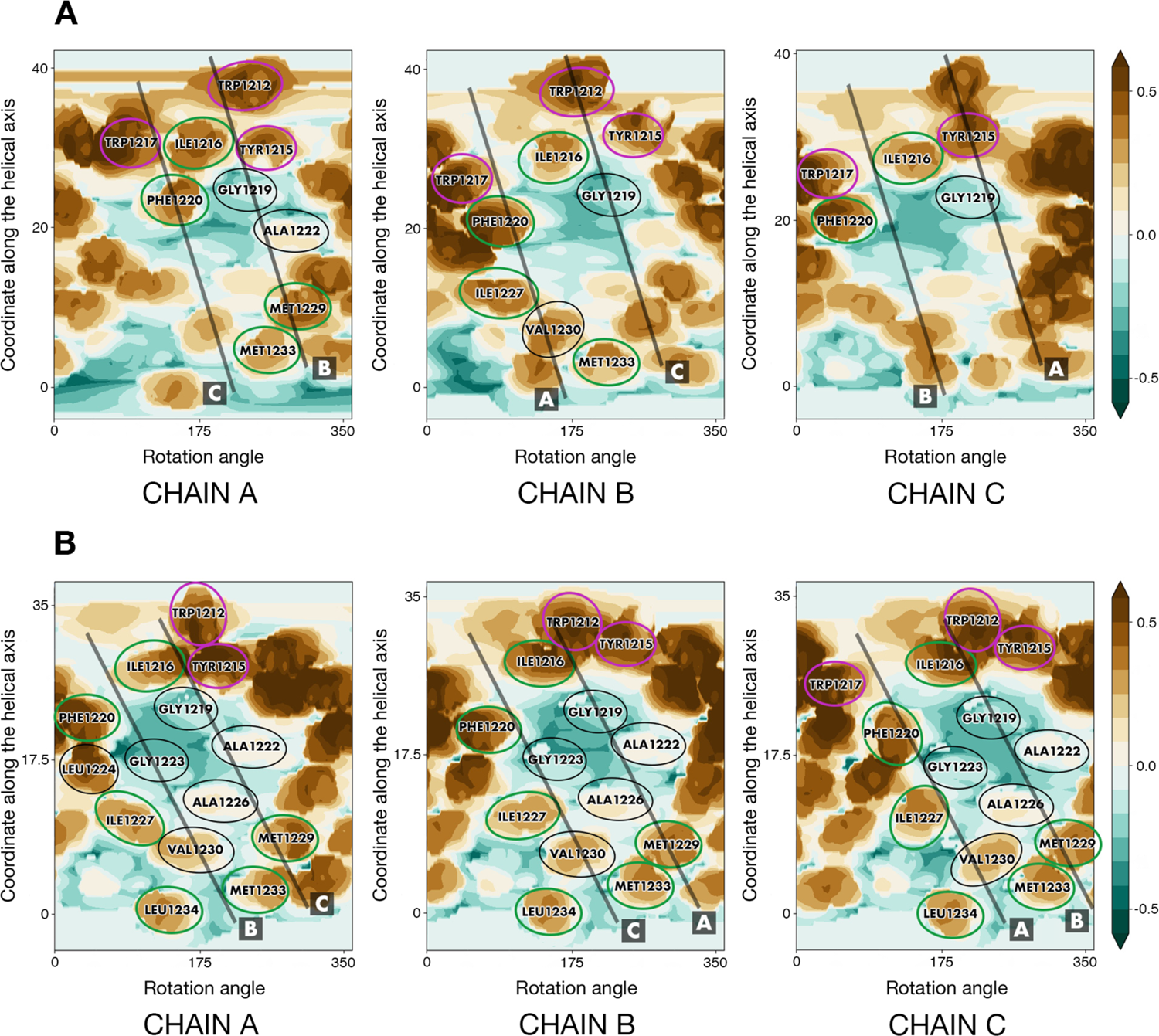
Helix/helix interfaces in *S_TNFR1* and *S_OPT*. (A) Helix/helix interfaces in *S_TNFR1* before simulation. (B) Helix/helix interfaces in *S_OPT* upon building and energy minimisation. Identical positions are encircled in purple, conservative and semi-conservative residues are encircled in green, and non-conservative residues present on the helix/helix interface are encircled in black. For each chain, projections of the other two helices are shown as semi-transparent black lines and designated by letters A to C. For other details see legend to Fig. 2B.

**Figure 5.**
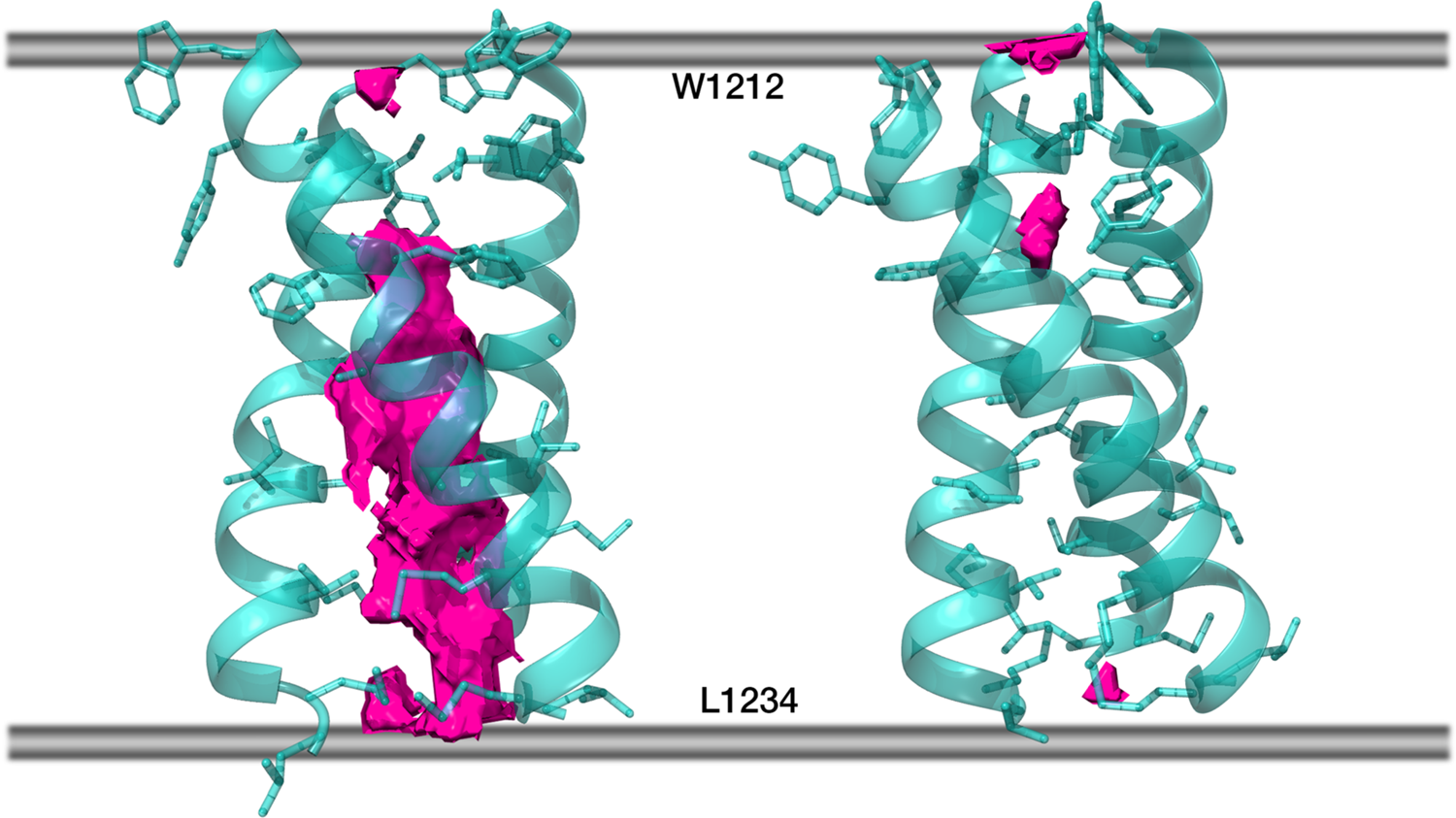
Free volume in the lumen of *S_TNFR1* (left) and in *S_OPT* (right). Protein chains are partially transparent and are shown in cartoon representation, residues within each chain facing either of the remaining two chains are shown in stick representation. Free volume is rendered as solid pink blocks (averaging over 200 frames not applied), while the boundaries of the hydrophobic stratum of the membrane are schematically shown as grey lines.

In *S_OPT* the distances between the axes of monomers range from 7.8 Å to 10 Å and the angles between helical axes equal 41±2°. Consistently found on the helix/helix interfaces were W1212, I1216, G1219, F1220, A1222, G1223, A1226, I1127, M1229, V1230, M1233 and L1234. Many of these positions (W1212, I1216, F1220, I1227, M1229, M1233 and L1234) can be considered at the very least semi-conservative within genus *Betacoronavirus*, which indicates their functional relevance to a protein encoded by a viral genome with a mutation rate characteristic of ssRNA viruses. Seeing that it was arranged as predicted and contained a remarkable number of (semi-)conserved residues on helix/ helix interfaces, a feature often observed in α-helical TMDs (Rees et al., 1989; Donnelly et al., 1993), *S_OPT* was selected to further test its stability.

### Model S_OPT is highly stable in a lipid bilayer: results of MD simulations

For the entire duration of the MD trajectories the model *S_OPT* remained stable: helix/ helix interfaces were undisrupted by lipids and retained a high degree of similarity to the model’s initial state (Fig. S3 A-C). In one of the trajectories, one of the helices moved slightly away from the original interface in its N-terminal portion (W1212/Y1215). Downstream therefrom, however, this chain contributed to the trimer interface as much as the other two.

The model trimer, which had initially been oriented along the normal to the membrane plane with residues 1212 to 1234 spanning the bilayer, demonstrated a tendency to remain in the described position, buried in the membrane and is slightly tilted relative to the normal to the bilayer plane. Indeed, the centre of mass of *S_OPT*’s residues 1212 to 1234 had a Z-coordinate corresponding to the midpoint between leaflets (Fig. S4 A, B), while α_ax_, the angle between the trimer principal axis and the normal to the bilayer, equalled 14°±7° and 12°±5° in trajectories *S_OPT run1* and *S_OPT run2*, respectively. The number of intermolecular P/P and P/L contacts did not vary significantly over the course of the MD simulations, indicating that extent of their exposure to the lipid environment did not alter (Table S2). Likewise, evaluation of FV in the lumen of the model TMD over the course of the MD trajectories indicates that it remains tightly packed and, whenever unoccupied volume can be detected, the trimer tends to revert back to a tightly packed state (Fig. 6, Fig S4 C), while the RMSD oscillations are consistent therewith (Fig. S4 D).

**Figure 6.**
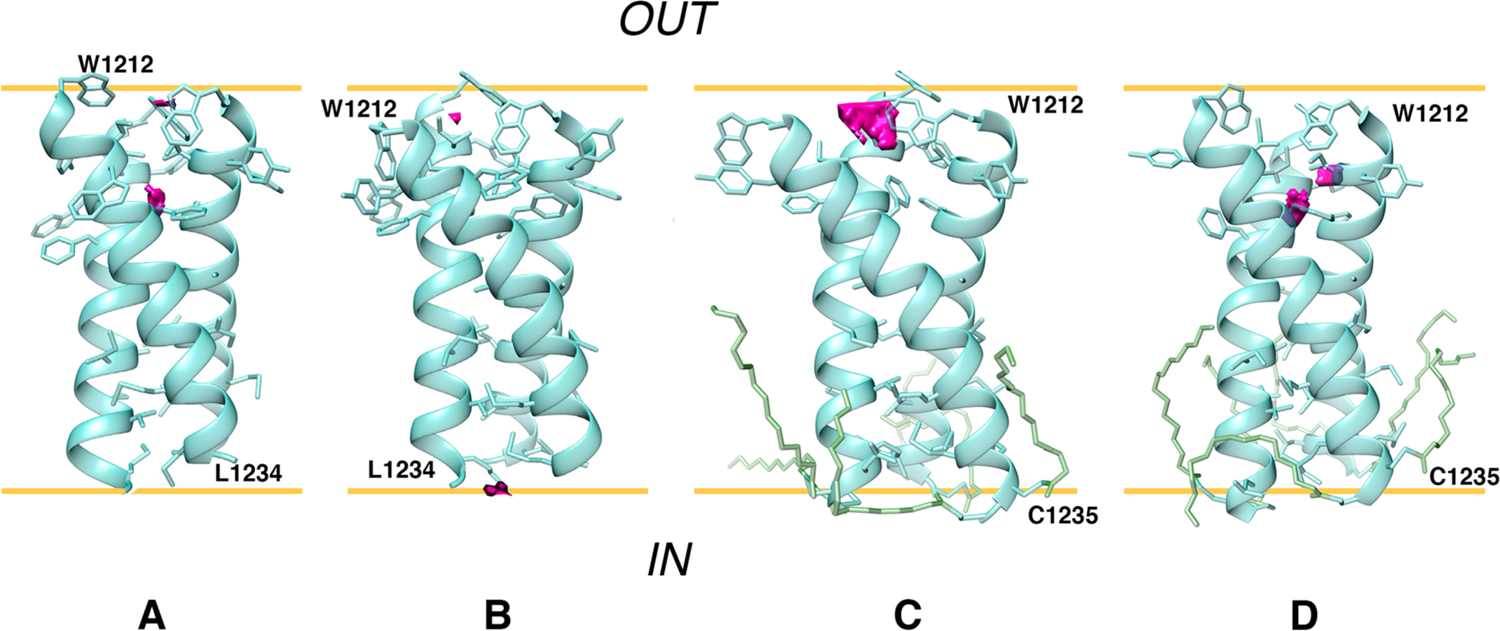
Visualisation of free volume in unpalmitoylated and palmitoylated *S_OPT*. Free volume in the lumen of the model trimers is rendered as pink blocks (averaging over 200 frames not applied) and shown at 1 μs of MD trajectories *S_OPT* run1 (A), *S_OPT* run2 (B), *S_OPT-PLM* run1 (C) and *S_OPT-PLM* run2 (D). 3D models are shown in cartoon/stick representation; palmitoyl chains are shown in stick representation where present. Approximations of POPC carbonyl/ester oxygen planes are shown as yellow lines. The inside of the virion and the external environment are designated as “IN” and “OUT”, respectively.

### Dynamic properties of the template TNRF-1 TMD and other S-TMD models

#### TNFR1_TMD

Over the course of a 500 ns MD simulation the angle α_ax_, amounted to 15±6° (as opposed to 12° at 0ns). Two stages can be clearly identified in the trajectory (Fig. S5). At first the RMSD of the backbone atoms of the TMD varies very little, until, at ∼238 ns, it begins to grow. Similarly, estimated FV in the TMD lumen seems to virtually disappear, resulting in tight packing, however, at around 250 ns it also starts rapidly increasing. At approximately the same moment side chains constituting the helix/helix interface in *TNFR1_TMD* start making an increasingly higher number of contacts with POPC molecules and gradually fewer contacts with each other, indicating that the helix/helix interface was disrupted by lipids. We therefore conclude that *TNFR1_TMD* was not stable in a model POPC bilayer, possibly partially due to an insufficiently long hydrophobic TM part, as, experimentally, the NMR structure was observed in DMPC/DH_6_PC bicelles and could thus be poorly adapted to a POPC bilayer. Indeed, immediately downstream of the proposed TMD is a flurry of polar and charged residues that require a hydrophilic environment.

#### Other S-TMD structures

Three models based on the recently published NMR TM trimer (PDB ID 7LC8) were tested (Table 1): the trimer structure verbatim (trajectory *S_NMR*), the same structure with short disordered fragments added upstream and downstream to stabilise the helices and minimise the “disordering” of our fragment of interest (trajectory *S_NMR+*) and 7LC8 with the sequence of the wild type spike protein (trajectory *S_NMR-WT*). The angle α_ax_ amounted to 27°±11°, 12°±6° and 21°±8° for *S_NMR*, *S_NMR+* and *S_NMR-WT*, respectively. For all the three trimers, we analysed the number of contacts between the residues constituting the Leu/Ile zipper proposed by the authors of the experimental study (I1221, I1225, L1229 and L1233) and lipids, as well as contacts these residues make with their counterparts in neighbouring helices. It appears that the number of contacts between Leu/Ile zipper residues within the trimer decreases over time, whereas the number of contacts these residues make with lipids grows. Thus, larger portions of the side chains became exposed to lipids, indicating that the helix/helix interfaces were disarrayed by POPC molecules, and the Leu/Ile zipper which had been observed experimentally in DMPC/DH_6_PC bicelles was destabilised, no longer holding the trimer together. Residues that had been initially tucked away into the trimer’s lumen become increasingly more exposed to the lipid environment (Fig. S6 A, B). In line with this is the estimated FV (Fig. S6 D, E) in the lumen of each of the three trimers examined. In each trajectory it dramatically fluctuates over time, but the trimer fails to revert back to a relatively tightly packed state it was in at the outset of the simulation with no lipids on the helix/helix interface. RMSD values (Fig. S6 C) also indicate that initial configuration lifetimes were short for this group of models. The structure experimentally observed in bicelles and its derivatives were thus not stable in a model POPC bilayer.

Both mutants containing substitutions present in the NMR model, *S_OPT-LC* and *S_OPT-LS*, remained tightly packed, while the angle α_ax_ equalled 13°±7° and 9°±4° for *S_OPT-LC* and *S_OPT-LS*, respectively. The behaviour of these trimers did not indicate significant destabilisation compared to their precursor with the wild type amino acid sequence, as evidenced by the RMSD and FV dynamics (Fig. S6 F, G), as well as by the numbers of intermolecular P/P and P/L contacts (Table S2). We thus observed no significant impact of the substitutions present in the experimentally observed model trimer on the organisation of helix/helix interfaces nor on the stability of the model trimer *S_OPT*. Therefore, instability in a lipid bilayer of the NMR model of S-TMD with 4 mutations in the sequence would appear to stem from other factors than these modifications. The reason is most likely to lie in the poor adaptation of the NMR-derived template (e.g., mutual packing of helices in a trimer) to the POPC bilayer. However, as the mutations were introduced into a pre-existing model trimer, one cannot rule out the possibility that they could affect TMD trimerization during assembly in the membrane.

The S-TMD model built by Casalino et al. (2020) has completely different helix/helix interfaces (Fig. S3 D), redolent of a canonical water-soluble coiled-coil, with the most voluminous hydrophobic residues tucked inwards facing the trimer lumen. This approach does not take into account the known principles of TM helix packing. We did not explore this model in further detail. Meanwhile, both S-TMD models created by Woo et al. (2020) have helix/helix interfaces very similar to those in *S_OPT* (Fig. S3 E, F) and dASA of 2450 Å^2^ / 2406 Å^2^. RMSD for the backbone atoms in residues 1212 to 1234 in these two trimers was 2.1 Å and 2.5 Å as compared to *S_OPT.* We tested them for stability in a POPC bilayer using the same fragment as in the case of *S_OPT* (1208-1238), and their performance was comparable to that of our model insofar as parameters like FV in trimer lumen were similar, not exceeding 10 Å^3^, and consistent throughout the MD simulations. Our model did somewhat better as far as RMSD goes, however: 1.3±0.2 Å and 1.2±0.3 Å for *S_OPT* as opposed to 2.9±0.7 Å and 2.4±0.2 Å for the two gp41-based models. Finally, model *S_OPT* is better packed based on dASA values (see above). These data suggest our model’s higher propensity for stability.

### Palmitoylation of S_OPT preserves its unique stability in a model membrane

Over the course of MD simulations both palmitoylated trimers retained their stability (Fig. S4 D), and their helix/helix interfaces were mainly similar to those in *S_OPT*. The angle α_ax_ for *S_OPT-PLM* amounted to 19°±7° in run1 and 16°±7° in run2. Whenever FV could be observed in a trimer’s lumen, the protein reverted to an extremely tightly packed state (Fig. S4 C), and the number of P/P and P/L intermolecular contacts also remained consistent over the course of the MD simulations (Table S2). The overall stability of the helix/helix interface appears to be comparable in palmitoylated and unpalmitoylated trimers, as is evidenced by RMSF calculations for each residue within the TMD (Fig. S4 E). The input of each residue into protein/protein contacts holding the trimer together is also uniform across all replicas calculated for *S_OPT* and *S_OPT-PLM*. The only deviation of relative magnitude observed in palmitoylated models is a greater input on the part of W1217 (Fig. S4 F).

Apart from preserving the trimer stability, the introduction of palmitoyl modifications does not cause significant perturbation of the lipid environment as compared to the non-palmitoylated trimer. Thus, the analysis of order parameter of lipid tails (S_CD_) reveals similar mosaic patterns in both cases, which comprise ordered and disordered bilayer regions resulting from the presence of the model TMD spanning the membrane (Fig. S7). Particularly, palmitoyl modifications do not promote any specific ordering of POPC lipids in the vicinity of S-TMD with a small tendency towards a decrease in corresponding S_CD_ values. These observations are not at variance with data on quasi-harmonic (QH) entropy (Fig. S4 G) evaluated for each lipid molecule in the system. Like in the case of a pure POPC bilayer used as a reference, all systems with either unmodified or modified S-TMD trimer yielded a near-normal distribution of lipid entropies. TΔS as compared to pure POPC averaged −0.32±1.19 kcal/mol and −0.36±1.30 kcal/mol per lipid for unpalmitoylated and palmitoylated trimers, respectively, indicating minor perturbation of the lipid bilayer similar for modified and unmodified S-TMD.

## DISCUSSION

Ways in which the machinery effectuating viral fusion operates are impossible to fathom without taking into account the role of the membranes, both of the host cell membrane and the viral envelope. This is due to the fact that the membrane is a major factor predetermining the structural and dynamic aspects of the behaviour of the key players involved in fusion, these being the TMD as well as, at later stages, amphiphilic domains like HR1 and HR2. The latter only become employed at a certain point, but their contribution is of great importance. Furthermore, it is imperative to bear in mind that the listed fragments of the spike protein impact each other, enabling the virus to completely fulfil its potential in fusing the membranes.

When evoking the importance of understanding the molecular aspects of TMD organisation and functioning, one shall have to think beyond its obvious role as anchor which ensures the correct positioning of the spike protein in the viral envelope. Not less crucial are the subtle features of its structure and dynamics, which presumably enable the spike to effectuate conformational rearrangements varying in scale from local to global and determining the transition from the pre-fusion to post-fusion state. For example, SARS-CoV spike, highly homologous to that in SARS-CoV-2 in the S2 subunit, showed much lower fusion activity upon introduction of point mutations in the aromatic-rich cluster. Corver et al. (2009) showed that, when two out of three tryptophan residues in this cluster (corresponding to W1212, W1214 and W1217 in SARS-CoV-2 spike) were substituted with phenylalanines, this resulted in a lack of viral entry for SARS pseudo-particles, while replacement of only one of these Trp residues resulted in a residual level of entry. Furthermore, substitution of polar residues Thr1238 and Ser1239 located C-terminally to Ala led to a significantly lower level of acylation of the spike protein (Panina et al., 2022). Acylation itself or the absence thereof had been demonstrated to bear an effect on the infectivity of both SARS-CoV (Petit et al., 2007) and SARS-CoV-2 (Mesquita et al., 2021) following substitutions of Ala for Cys residues 1235 and 1236 in the first cysteine cluster downstream of TMD (SARS-CoV-2 numbering). More or less similar effects have been reported in SARS-CoV spike for point mutations G1201K (G1219 in SARS-CoV-2 spike) and V1210K (V1228 in SARS-CoV-2 spike) (Corver et al., 2009). Finally, it has been experimentally demonstrated (Arbely et al, 2006) that the GxxxG motif in SARS-CoV spike (corresponding to 1219-GFIAG-1223 in SARS-CoV-2 spike) TMD is crucial to its trimerization, as substituting the first one of these glycines or both of them resulted in a dramatic reduction of oligomerization. (It is of note that these residues lie on the helix-helix interface in our model *S_OPT*.) To a considerable extent this effect was observed at the level of the entire spike protein, not just isolated TMD. Taken together, these results demonstrate that the functioning of such a complex machinery as the entire virion is sensitive to rather subtle modifications in TMD, which would presumably not be the case if TMD only served as a simple membrane anchor. Besides, it would seem highly likely that SARS-CoV-2’s S-TMD is organised following the same principles as the one in SARS-CoV, insofar as their TMDs are almost identical.

In order to propose a reliable configuration of TMD trimer corroborated by existing experimental observations, a comprehensive modelling framework was employed here with careful consideration of TMD boundaries and helix-helix association interfaces. Particularly, for the first time, homology modelling was done adopting a combined algorithm of template selection, which relied on the search of TM α-helical homotrimer structures followed by singling out ones wherein the physicochemical properties of individual monomers bore maximum resemblance to those in S-TMD. Such resemblance was recognised as similarity between patterns of residues in their sequences possessing comparable physical and chemical properties, as well as similarity between the distributions of hydrophilic, hydrophobic and variability-prone properties on helix surface, also known as “dynamic molecular portraits” (see section “*S_OPT*: a tightly packed TMD model obtained via iterative refinement”). Furthermore, the initial template-based models were additionally adjusted taking into account the results of PREDDIMER (Polyansky et al., 2014) prediction of helix-helix packing in order to optimally align the target protein and the template. This was done due to the lack of homology of their sequences. Importantly, microsecond-long MD simulations in a model POPC membrane helped to improve the stability of the model via its iterative fine-tuning. This step was necessary, seeing that direct mounting of the S-TMD sequence even onto the best-suited template led, as we observed, to unstable behaviour of the trimer in a water-lipid environment. This was due to the presence of hollow spaces (FV) in the trimer lumen, translating as suboptimal α-helix packing. These imperfections were identified in the course of MD and rectified manually, taking into account PREDDIMER results. The proposed model, herein referred to as *S_OPT*, eventually demonstrated a very high level of stability in independent runs of microsecond-range MD simulations. Interestingly, in most cases, models based on structures observed experimentally in membrane-like environments (detergent micelles, bicelles, etc.) do not demonstrate such stability in a POPC bilayer.

As opposed to tools for computational prediction of transmembrane dimer structure (Polyansky et al., 2014; Cao et al., 2017) that have proven to be adequate as a result of comparison against existing experimental models (including ones obtained post factum), efficacious and generalised methods of transmembrane trimer modelling have not yet been developed. Targeted sporadic attempts at such modelling have been made using regular symmetrical search, simulation annealing and energy minimization (Arbely et al., 2006) for SARS-COV spike, using PREDDIMER for the prediction of association interfaces for individual helical pairs and their combination for influenza hemagglutinins (Kordyukova et al., 2011), or via multiscale simulations of self-assembly for individual TMDs of T-cell receptor (Sharma et al., 2013). The present study, on the other hand, demonstrates a consistent framework combining reliable prediction tools and MD simulations, which enables one to model noncovalently associated trimers of TM helices following a systematised procedure.

The proposed model was also shown to be stable in microsecond-range MD simulations, when two cysteine residues located immediately downstream of the TMD are palmitoylated. The modifications do not display significant constraining or “freezing” of POPC lipids in the vicinity of the trimer, while general membrane perturbation effect is similar to that of the non-palmitoylated trimer, as is evidenced by the analysis of lipid order parameters and their configurational entropies. However, due to limitations of the MD simulations performed such as the single-component POPC bilayer and elevated temperature, fine-tuning of the lipid environment due to palmitoylation as well as its functional interpretation is not fully accessible in our modelling framework and should be further investigated. Apparently, such effects can be observed in lipid bilayers more complex in composition than POPC; some recent experimental data demonstrate the importance of this issue (Vilmen et al., 2021)

Seeing that the proposed TMD model has been created *in silico* and does not directly rely on experimental structural data, questioning its accuracy and level of similarity to other models would not be unjustified. As far as accuracy is concerned, only one structure of the TMD has been reported, which was obtained by NMR spectroscopy in lipid-detergent bicelles (Fu and Chou, 2021). Helix/helix interfaces in this structure differ from those proposed in the present work (see section “Dynamic properties of the template TNRF-1 TMD and other S-TMD models”). However, the NMR model has a number of features prompting one to wonder whether it corresponds to the native state of the TMD. Most importantly, its amino acid sequence contains four substitutions and one deletion, which could dramatically impact helix packing. Furthermore, bicelles as a membrane-mimicking environment might significantly differ from a lipid bilayer, especially in the vicinity of the TMD boundaries. In the study in question, the N-terminal portion of the peptide under investigation (corresponding to the aromatic-rich cluster) was not part of the TMD, and its structure was not resolved, while the segment spanning the membrane was short, only containing 16 residues (from L1218 to L1234). The authors observed a disordered fragment inclusive of residues 1212 to 1217, whereas this region in SARS-CoV spike had previously been shown to assume a conformation redolent of helical (Mahajan and Bhattacharjya, 2015) and to have an affinity for water/lipid interfaces. The omission of a cluster of aromatic residues capable of strongly interacting with each other and with the membrane environment could seriously affect the geometry of the entire TMD model. Our calculations indicate that the NMR model of the TMD does not remain compact in a POPC bilayer, to the likes of which it is probably poorly adapted. However, this circumstance cannot be regarded as proof of the conclusion drawn above. In our model, this fragment is close to the water/lipid interface, in agreement with experimental data indicating that this is where it tends to localise. Its propensity for existing in the helical conformation (in which it is rendered in our model) means that it could form one entity with residues located downstream, also in the helical conformation.

The question of the structural/dynamic organisation of the aromatic-rich region downstream of HR2, sometimes referred to as the pre-transmembrane domain (Guillén et al., 2007), is very important because it serves as a kind of “hinge” that provides large-scale conformational transformations accompanying the transition from the pre- to the post-fusion state (Walls et al., 2017; White et al., 2008). Given the trimeric nature of this fragment and its richness in aromatic and positively charged residues, which can strongly interact with each other in different combinations, this domain most likely has high conformational plasticity. It is this quality that would allow it to perform the crucial mechanistic role in the process of merging the membranes of the virus and the target cell. Hence attributing a single precise structure to such domains would not seem to make a lot of sense. One could draw an analogy with known membrane proteins like receptor tyrosine kinases (RTK), such regions on the membrane-water interface can adopt various conformations, depending on the external forces coming into play at different moments during the effectuation of their function, as well as on their exposure to varying environmental conditions (Bocharov et al., 2018). A key role in such events can be played by proline residues, the peptide bond in which can adopt both cis- and trans-configurations, providing regulation of the inclination of the N-terminal portion of the TM helix and the adjacent extracellular domain with respect to the membrane (Crnjar et al., 2019). Note that in the spike proteins, the membrane-proximal proline residue (Pro1213 in the spike of SARS-CoV-2) is highly conservative. There is a distinct sublineage of viruses (Yu et al., 2020), whereof spike proteins sport a lower than 28% identity to other coronaviral spikes, in which this position is occupied by alanine. However, as far as the apparent importance of proline in this position in other lineages goes, further experimental studies, in particular, using NMR spectroscopy, could shed light on its contribution to the fusion process.

Our model of the S-TMD differs from other models thereof created as part of large-scale projects on full-length spike modelling such as one based on a water-soluble coiled-coil template (Casalino et al., 2020). Interestingly, though, a model of the TMD built using HIV’s gp41 protein (Woo et al., 2020) is strikingly similar to ours. They proposed two versions of the TMD to use in a range of their full-length spike models, and both of them turned out to possess helix-helix interfaces very similar to those in *S_OPT*, although the gp-41-based model is still less stable in a POPC membrane than the one proposed in the present work. Furthermore, the authors only palmitoylated Cys1236, but not Cys1235, identifying it as solvent-exposed. However, it is known that replacement of both these cysteines with alanine leads to a dramatic drop in the level of acylation, 80% for the spike protein of SARS-CoV2 (Mesquita et al., 2021) and 56% for the spike of SARS-CoV (Petit et al., 2007). While the input of palmitoylation of these two cysteines separately has not been verified, one could speculate that both of them might be solvent-exposed, as one ninth of all cysteine residues in all palmitoylation clusters within the endodomain is less likely to account for 80% of palmitoylation compared to two-ninths thereof. Our model offers both Cys1235 and Cys1236 available for palmitoylation and positioned in such a way that, if palmitoylated, this does not affect the stability of the TMD trimer. Nevertheless, it is interesting that, as far as helix/helix interfaces within the TMD are concerned, our model and the one based on gp-41 demonstrate a great level of convergence, having been obtained using completely different approaches. Like spike, gp-41 is a class I fusion protein, and one cannot help wondering whether the convergence of these two models is due to features peculiar to viral fusion protein TMDs in addition to general principles of transmembrane α-helix packing.

Equally of note is the fact that the proposed trimer model can provide a direct structural explanation of the effect of some of the TM mutations mentioned above, as well as the role of the GxxxG motif, since, in our model, the residues in question belong to the interfaces of helix association (Figure S4 F) and their substitution may modulate stability and configuration of the trimer.

Importantly, the *S_OPT* model has already been put to practical use to verify the MHP compatibility between the spike protein and hDHHC20, the enzyme conducting palmitoylation. In our model the hydrophilic patch, formed by T1231, S1239 and the highly conserved throughout all geni of coronaviruses T1238, is solvent-exposed. T1238 and S1239 were shown to be crucial to acylation via mutagenesis, and it is thus hypothesised that the hydrophilic patch is part of the docking site for hDHHC20, which has a hydrophilic patch of its own next to where the fatty acid chain is harboured (Panina et al., 2022). Our model therefore does not contradict experimental data on the accessibility of this potential docking site to the enzyme effectuating palmitoylation.

The purpose of the present study was not to model the way TMD operates, for instance, during the transition from the pre-fusion to the post-fusion state. This is the subject of further exploration, which will require gradual complication of the system, i.e., the addition of regions adjacent to the TMD. Most notably one will have to consider domains HR1 and HR2 consisting of amphiphilic α-helices, major factors in spike refolding during fusion that eventually form the so-called “trimer of hairpins” made up by 6 helices and stabilising the post-fusion state. It has already been demonstrated that HR regions possess affinity for the membrane apart from affinity for each other, HR1 much more so than HR2 (Chiliveri et al., 2021). It is thus possible and has indeed been speculated that such interactions facilitate viral fusion. Our model could be useful in the exploration of the behaviour of these membrane-proximal fragments of the spike protein involved in fusion, most notably HR2, with recourse to computational tools similar to those employed in the present study.

## METHODS

### Assessment of TMD boundaries via sequence analysis and Monte Carlo simulations with an implicit membrane

Initial guesses regarding the boundaries of the TMD were made using a set of bioinformatics tools applied to the spike sequence. Based on the predictions made using TMSEG (Bernhofer et al., 2016) and TMPRED (Hofmann and Stoffel, 1993), as well as taking into account data on proposed TMD boundaries from UniProt entry P0DTC2 (The UniProt Consortium, 2021) and related publications, the peptide selected for more detailed assessment of the structure and membrane localization of the S-TMD corresponded to residues 1208 to 1239 (QYIK<u>WPWYIWLGFIAGLIAIVMVTIML</u>CCMTS). It was then examined via MC simulations with an implicit membrane model (Nolde et al., 2000). The starting structure of the TMD was constructed in the α-helical conformation (underlined in the aforementioned sequence).

Peptide charges were assigned as if it remained at pH 7. At the beginning of MC calculations with an implicit membrane model, the peptide was arbitrarily placed outside the hydrophobic layer, or the so-called “hydrophobic slab”. To change the orientation of the peptide with respect to the slab during an MC simulation, a fragment of 21 dummy residues was attached to its N-terminus. These “virtual” residues did not contribute to the energy of the system. The first atom of the N-terminal dummy residue was always placed at the centre of the slab and assigned coordinates (0,0,0).

The conformational space of the peptide was explored via unrestrained MC search in dihedral angles space using the modified FANTOM program (von Freyberg and Braun, 1991) and our implicit solvation model for the membrane-mimic heterogeneous water/ cyclohexane environment (Nolde et al., 2000). The all-atom potential energy function was expressed as *E_total_* = *E_ECEPP/2_* + *E_solv._*, where the term *E_ECEPP/2_* includes van der Waals, torsion, electrostatic and H-bonding contributions (Némethy et al., 1983). *E_solv._* is the solvation energy calculated as follows:

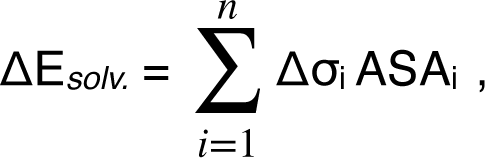

where Δσ_i_ and ASA_i_ are the atomic solvation parameter (ASP) and the solvent-accessible surface area (ASA) for atom i, respectively. ASP values for gas/water and gas/cyclohexane transfer were taken from (Efremov et al., 1999). To cross the energy barriers between local minima, the adaptive-temperature schedule protocol (von Freyberg and Braun, 1991) was employed. At each MC-step the structures were subjected to 50 to 100 steps of conjugate gradient minimization. The ω dihedral angles were fixed (except those in dummy residues), and a spherical cutoff of 20 Å for non-bonded interactions was employed. Long-range electrostatic interactions were treated with distance-dependent dielectric permeability calculated as ɛ=4×r. The half-width of the membrane interface region and the slab thickness were 1.5 and 30 Å, respectively.

Before the simulations, starting structures were subjected to 100-200 steps of unconstrained conjugate gradients minimization. For each peptide, several consequent MC runs were carried out. At the initial stages of the MC-protocol, large structural changes were allowed via the sampling of several dihedrals, after which more detailed conformational search with only few varied dihedrals was performed. The starting structure for each MC run was the lowest-energy one obtained during the previous run. The set of resulting structures was analysed using the following parameters: total energy, Z-coordinate of the centre of mass of the TMD, secondary structure, depth of immersion of residues in the membrane, tilt angles of the helical segment with respect to the normal to the membrane plane (axis Z), hydrogen (H) bonding. Analysis of MC data was done using auxiliary programs specially written for this purpose. Along with other details of the computational protocol they are described elsewhere (Efremov et al., 1999; Nolde et al., 2000).

### Search for the template for homology modelling of S-TMD

#### Analysis of TM sequences in homotrimers of TMDs with known spatial structure

Information on such homotrimers - candidate templates for S-TMD modelling - was gleaned from the OPM database (Lomize et al., 2021). Sequences of these TM fragments were then compared against that of S-TMD in order to identify patterns of residues with physicochemical properties in the TM sequence sufficiently similar to those in the fragment of interest in the spike. Positions of the following types of residues were taken into account: charged, polar, hydrophobic, small and prolines (see Table S1).

### MHP mapping and helix/helix interface analysis

Molecular hydrophobicity potential (MHP) maps were calculated as described elsewhere (Efremov et al., 1992; Pyrkov et al., 2009). This was done for candidate templates and spike TM segment 1212 to 1234 built in an ideal α-helical conformation. Helix-helix dimerization interfaces were predicted using the PREDDIMER web server (https://preddimer.nmr.ru) (Polyansky et al., 2014). ASA values were calculated using DSSP (Kabsch and Sander, 1983). A residue was considered to be located on the interface if the difference between its ASA values in a single helix and in a helical oligomer was ≥ 25 Å^2^ (≥ 10 Å^2^ for glycine).

### Modelling of S-TMD

Template-based modelling of the S-TMD was carried out in MODELLER9.19 (Šali and Blundell, 1993). The initial model was built based on the NMR structure of the TMD of TNFR-1 (PDB ID 7K7A) (Zhao et al., 2020). Small unordered sequences (Table 1) were added upstream and downstream of the TMD also using Modeller 9.19, the final model eventually including residues 1208-1238. *S_OPT*, the refined and better packed model trimer, was built in Pymol 2.4.0 based on MD-derived dimerisation interfaces of tightly packed helices in the TNFR-1-based trimer (see Results). Tight clashes between residues were eliminated in UCSF Chimera 1.14 (Pettersen et al., 2004). In all cases, point mutations were introduced in MODELLER9.19.

Introducing palmitoyl modifications at C1235 and C1236, two different initial orientations of the short unordered peptide fragments downstream of L1234 were tested, along the TMD axis and parallel to the bilayer plane. In both cases all six palmitoyl chains were oriented along the normal to the bilayer plane. Topology files for palmitoylated TMD monomers were generated using in-house software.

### MD simulations

On all occasions the TMD fragment was aligned with the normal to the bilayer plane and was inserted in the system in such a manner as to span the membrane. MD simulations were performed using the GROMACS 2020.4 package (Abraham et al., 2015) and the CHARMM36 force field (MacKerell et al., 1998; Vanommeslaeghe et al., 2010; Best et al., 2012; Klauda et al., 2012; Huang et al., 2017). An integration time step of 2 fs was used and 3D periodic boundary conditions were imposed. The spherical cut-off function (12 Å) was used to truncate van der Waals interactions. Electrostatic interactions were treated using the particle mesh Ewald (PME) method (Essmann et al., 1995) (real space cutoff 12 and 1.2 Å grid with fourth-order spline interpolation). The TIP3P water model was used (Jorgensen et al., 1983), and Na^+^ and Cl^−^ ion parameters for counter ions were implemented. Simulations were performed at 325 K temperature and 1 bar pressure maintained using the V-rescale (Bussi et al., 2007) and the Parrinello–Rahman (Parrinello and Rahman, 1981) algorithms with 0.5 and 5.0 ps relaxation parameters, respectively, and a compressibility of 4.5 × 10^−5^ bar^−1^ for the barostat. The protein along with membrane lipids and solvent molecules were coupled separately. Semi-isotropic pressure coupling in the bilayer plane and along the membrane normal was used in the simulations. Before the production runs, all systems were minimised over 5000 steps using a conjugate gradients algorithm, followed by heating from 5K to 325 K over 50000 steps, during which internal coordinates of the protein and ligand heavy atoms were restrained. Production runs were simulated for 0.5 to 1.0 μs depending on the system (see Tables 1 and 2 for details). Bonds with an H atom were constrained via implementing LINCS (Hess et al., 1997).

### Data Analysis

#### MD trajectory analysis

All MD trajectories were processed using the *trjconv* utility from the GROMACS 2020.4 package to get the protein centred in the box, 3D periodic boundary conditions removed and to obtain an output frequency of 100 ps per frame. Coordinates were extracted using the *traj* utility, while RMSD and RMSF were calculated using the *rms* and *rmsf* utilities, respectively, all from the GROMACS package. Intermolecular contacts were calculated using PLATINUM (Pyrkov et al., 2009) and other in-house software (e.g., Krylov and Efremov, 2021). Molecular editing and graphics rendering were performed using PyMOL v.2.4.0 (*The PyMOL Molecular Graphics System, Schrödinger, LLC*) and UCSF Chimera package v. 1.14 (Pettersen et al., 2004).

#### Free volume calculation

For each trajectory frame the trimer and surrounding molecules were oriented along the axes of a local coordinate system to calculate the 3D distribution of free volume (FV) available to probe spheres with a radius of 1.4 Å (the size of a water molecule or a CH3/ CH2 group) within a regularly spaced rectangular mesh inside the trimer. Nodes of the mesh were considered free if their average occupancies over the analysed frame sequence were lower than 0.5.

A regularly spaced rectangular mesh was placed so that the X axis of the local coordinate system (CS) would pass through the centres of mesh faces parallel to the YOZ plane, while the coordinate system origin would be on the lower YOZ face centre. Local CS origin coincided with the centre of mass (CoM base) of CA atoms in S-TMD residues 1234 and 1235 (residues 233 and 234 for TNFR-1). Its X axis passed through the CoM of CA atoms in S-TMD residues 1211/1217 and 1212/1218 (depending on the model) or residues 211 and 212 in TNFR-1 (CoM top). The CA atom of residue 1234/233 in S-TMD/TNFR-1 chain A lay in the YOZ plane. The dimensions of the mesh were chosen empirically in order for the mesh to cover the TMD trimer lumen area (Fig. S8). Nodes of the mesh were positioned with an increment of 0.5 Å along all axes. For each trajectory frame, the protein and surrounding molecules were oriented as described above, and free volume (FV) values were calculated. Initially all mesh vertices had a weight of 0. During FV estimation each van-der-Waals (vdW) sphere associated with an atom was checked against nearby mesh vertices. All vertices lying inside a given vdW sphere were assigned weight equal to 1.

The data thus collected was used for the evaluation of the FV in the TMD trimer lumen, which were conducted as follows: all nodes where occupancy time was less than 50% were singled out, connected components of FV mesh nodes in 3D space were identified, and the largest component was selected. The node count of the largest component was then multiplied by the cell volume, and the size of its bounding box along the direction of the X axis was derived therefrom, providing estimated FV in the trimer lumen. FV fluctuations throughout MD trajectories were calculated in this manner applying a moving average over 200 frames.

#### Membrane response evaluation

Conformational entropy for lipids was evaluated using the quasi-harmonic (QH) approach employing mass-weighted covariance matrices in Cartesian coordinates derived via the *covar* utility from the GROMACS package. The resulting eigenvalues were processed to calculate QH entropies as described previously (Polyansky et al., 2012). For each system that underwent such analysis QH entropies were calculated per lipid molecule. Conformational entropy values were evaluated relative to those in the “pure” POPC bilayer, which was simulated for 1 µs using the same MD protocol as for other systems (see above). For entropy calculation, the first 0.25 µs of each trajectory were omitted to avoid artefacts related to system equilibration.

Carbon-deuterium order parameters (S_CD_) for the acyl chains in POPC molecules were calculated as described elsewhere (Douliez et al., 1998), while S_CD_ maps were created using in-house software. To this end, S_CD_ parameters for atoms in lipid acyl chains were calculated for all frames in the MD trajectory prepared with recourse to the *-fit rotxy+transxy* setting in the *trjconv* utility from the GROMACS 2020.4 package. Coordinates of the atoms were used to distribute S_CD_ values over a two-dimensional 50 × 50 units grid on the membrane plane equalling in size to the simulation cell dimensions. The data were averaged over the trajectory and used to plot the S_CD_ maps for each bilayer leaflet.

## DATA AVAILABILITY

The model of SARS-CoV-2 spike TMD proposed in the present study is available upon request in PDB.

## ACKNOWLEDGEMENTS

The project was supported by the Russian Science Foundation (grant 18-14-00375). Supercomputer calculations were performed within the framework of the HSE University Basic Research Program. Access to computational facilities of the Supercomputer Centre “Polytechnical” at the St. Petersburg Polytechnic University is gratefully appreciated.

## AUTHOR CONTRIBUTIONS

Conceptualisation, R.E.; Methodology, R.E., A.P., and E.A.; Investigation, Formal Analysis, Writing – Review & Editing, E.A., N.K., A.P., and R.E.; Software, N.K. and D.N.; Visualisation, E.A. and N.K.; Writing – Original Draft, E.A., A.P., and R.E.; Supervision, Project Administration, R.E.; Funding Acquisition, R.E.

## SUPPLEMENTARY INFORMATION

**Table S1.**
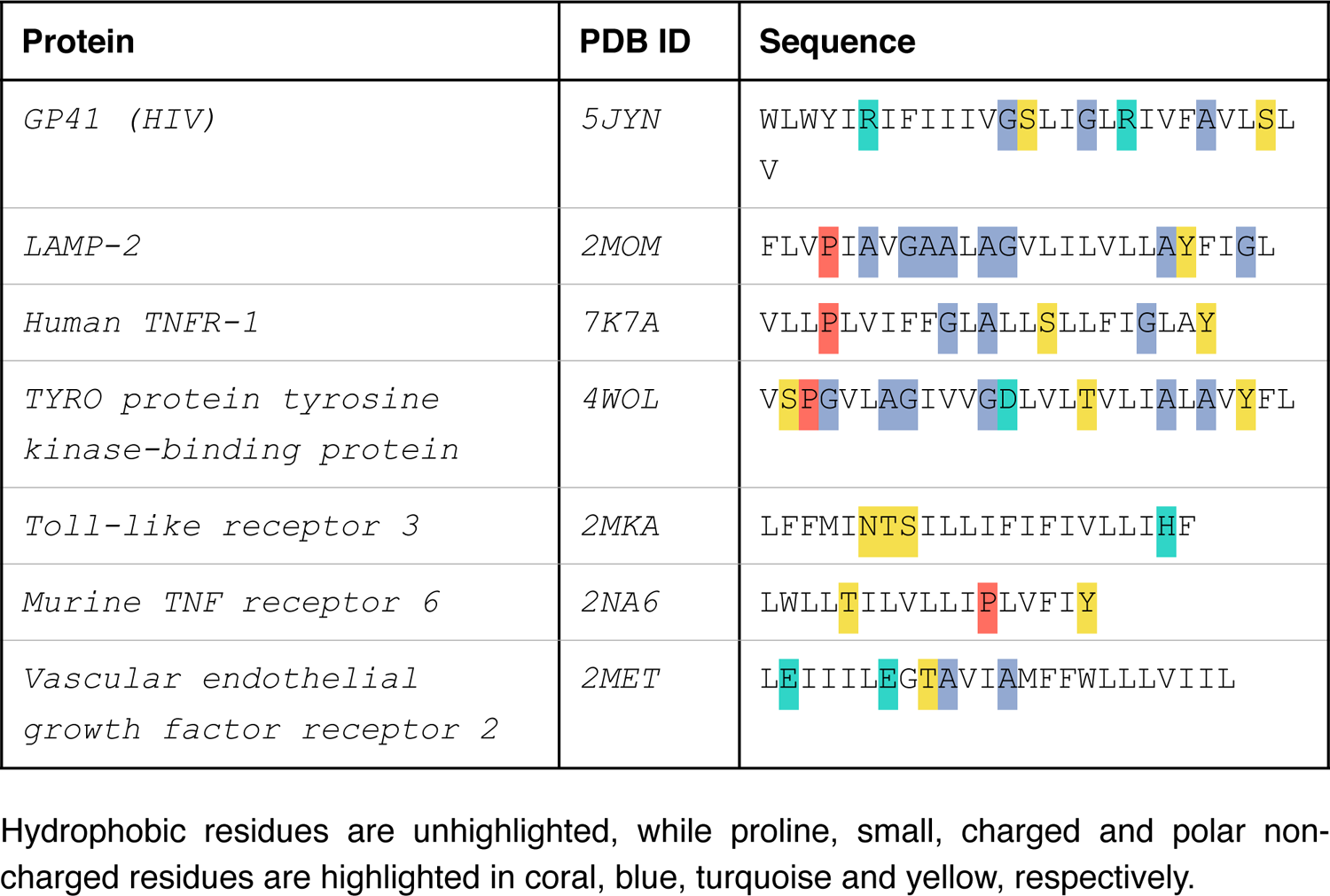
Amino acid sequences of TMDs for which structural data are available.

**Table S2.** Intermolecular protein-protein (P/P) and protein-lipid (P/L) contacts^1^ in *S_OPT* and its derivatives over the course of MD simulations.

| Trimer | P/P | P/L<sup>2</sup> |
| --- | --- | --- |
| *S_OPT run1* | 199±5 | 126±9 |
| *S_OPT run2* | 199±6 | 124±10 |
| *S_OPT-LC* | 191±6 | 130±9 |
| *S_OPT-LS* | 197±6 | 127±9 |
| *S_OPT-PLM run1* | 197±7 | 113±10 |
| *S_OPT-PLM run2* | 199±6 | 114±10 |

<sup>1</sup>: The number of pairs of atoms engaged in intermolecular contacts formed by the side chains of *S\_TMD*'s lumen-facing residues (1212, 1216, 1219, 1220, 1222, 1223, 1226, 1227, 1229, 1230, 1233, 1234) was calculated.
<sup>2</sup>: including protein-bound palmitoyl chains, when present.

## SUPPLEMENTARY FIGURE CAPTIONS

**Figure S1.**
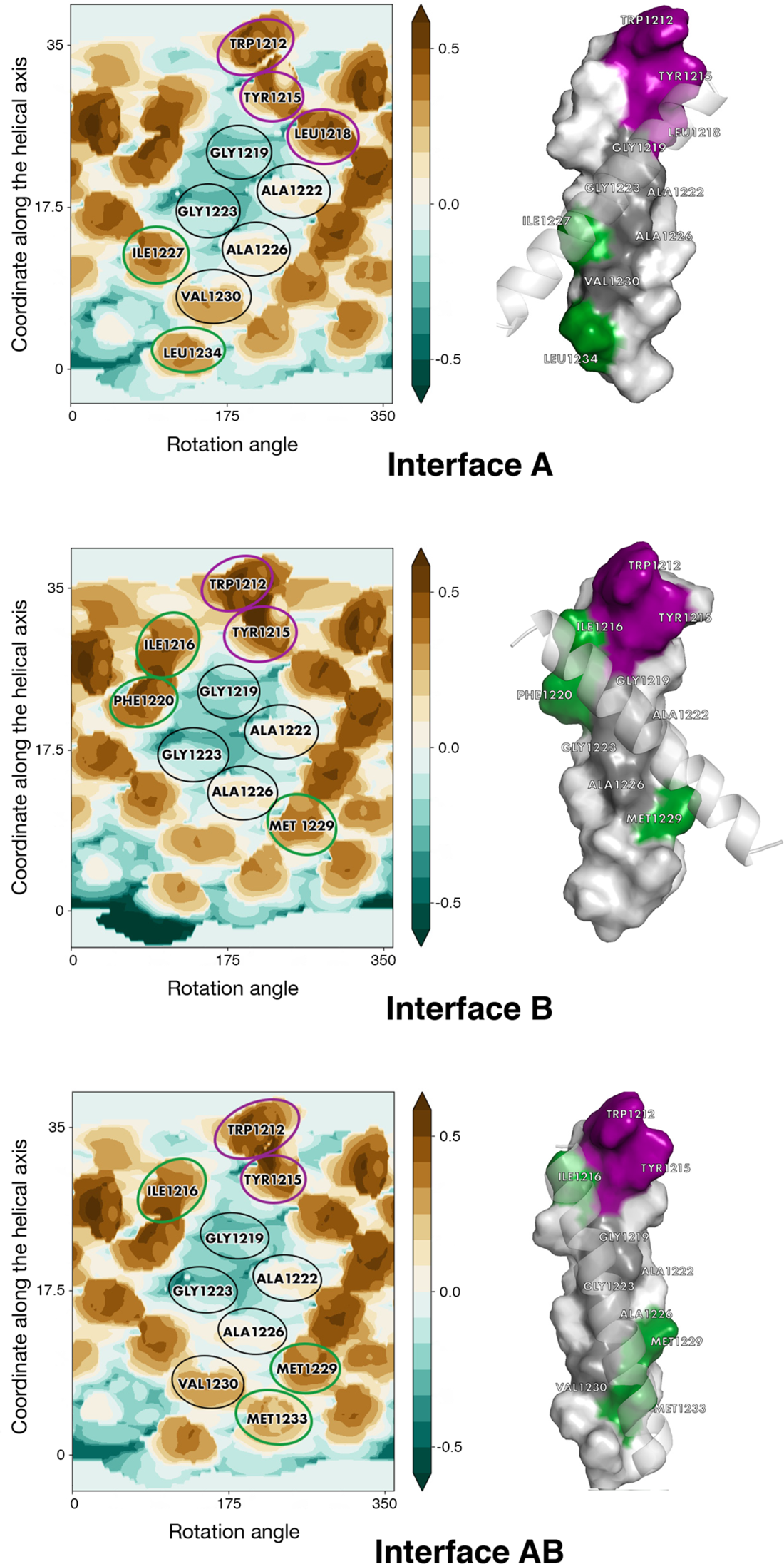
Dimerization interfaces predicted for S-TMD. Dimerization interfaces were predicted for an ideal helix with a sequence corresponding to S-TMD residues 1212-1234. In the MHP maps, identical positions are encircled in purple, conservative and semi-conservative residues are encircled in green, non-conservative residues present on the helix/helix interface are encircled in black. Cylindrical projection of the surface MHP distribution is used. Axis values correspond to the rotation angle around the helical axis and the distance along the latter, respectively. MHP scale (in logP octanol-1/water units) is presented on the right. The maps are coloured in accordance with the MHP values (Efremov et al., 1992), from teal (hydrophilic areas) to brown (hydrophobic ones). In the 3D models corresponding to each interface, identical and (semi-)conservative positions coloured in accordance with the same colour code as in the MHP maps, non-conservative residues located on the interface are coloured grey; surface representation of one of the helices is offered, while the second helix is shown in cartoon representation and is semi-transparent.

**Figure S2.**
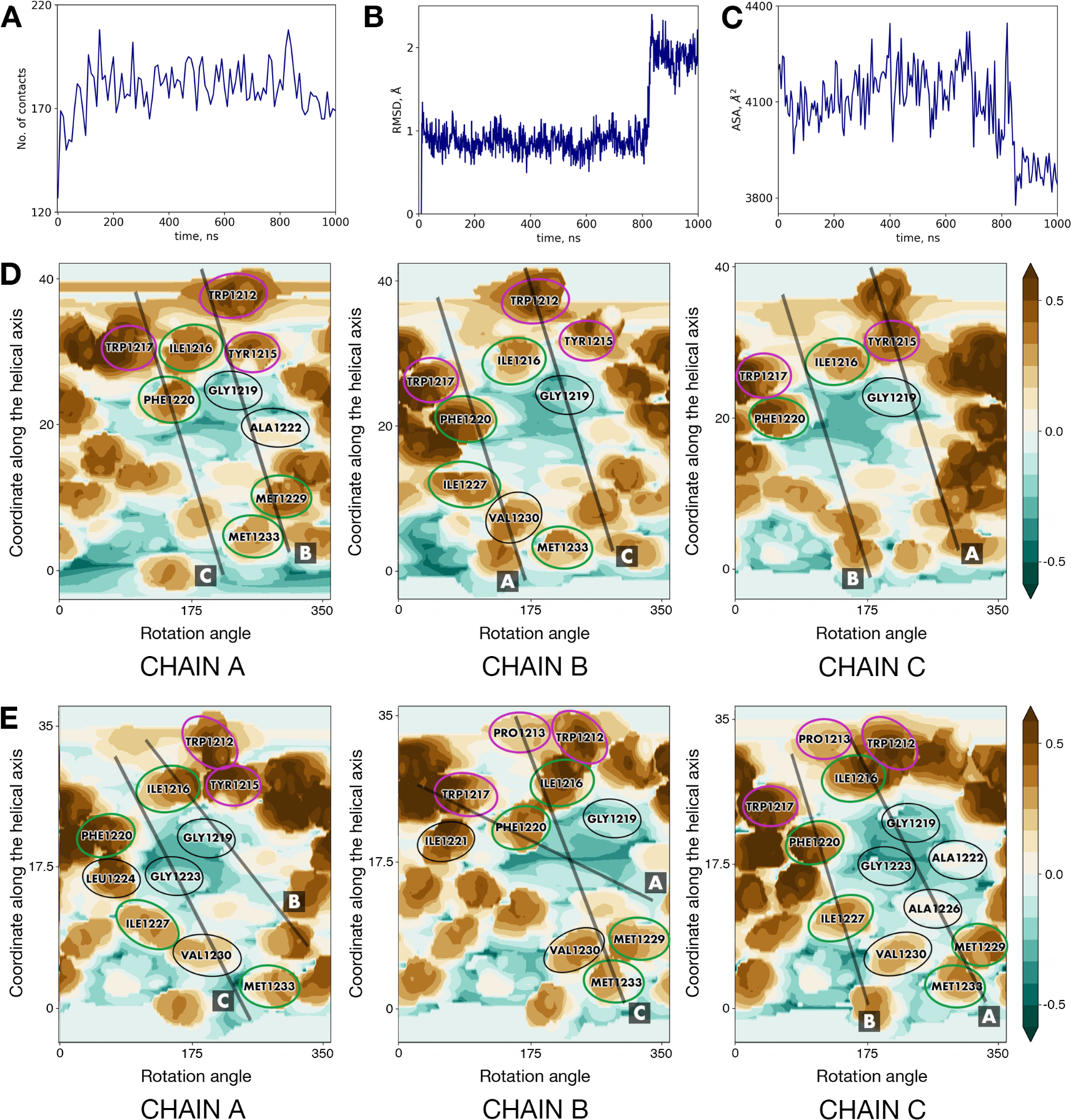
Performance of *S_TNFR1* in a POPC bilayer. (A) Protein/lipid contacts for the side chains in (originally) lumen-facing residues (1212, 1216, 1219, 1220, 1222, 1223, 1226, 1227, 1229, 1230, 1233) in *S_TNFR1*. (B) RMSD dynamics calculated for the backbone atoms of residues 1212-1234 in the two chains of *S_TNFR1* (A and C) that went on to form a highly stable dimer. The state registered at 9ns, the first frame when it was detected, was used as the reference frame for RMSD calculation. (C) Solvent-accessible surface area (ASA) for the same dimer made up by chains A and C. (D) Helix/helix interfaces in *S_TNFR1* at 0ns. (E) Helix/helix interfaces in *S_TNFR1* at 200ns. Identical positions are encircled in purple, conservative and semi-conservative residues are encircled in green, non-conservative residues present on the helix/helix interface are encircled in black. In the MHP map for each chain, schematic projections of the other two chains are shown as thick black lines and are designated by letters A to C. For other details see legend to Fig. S1.

**Figure S3.**
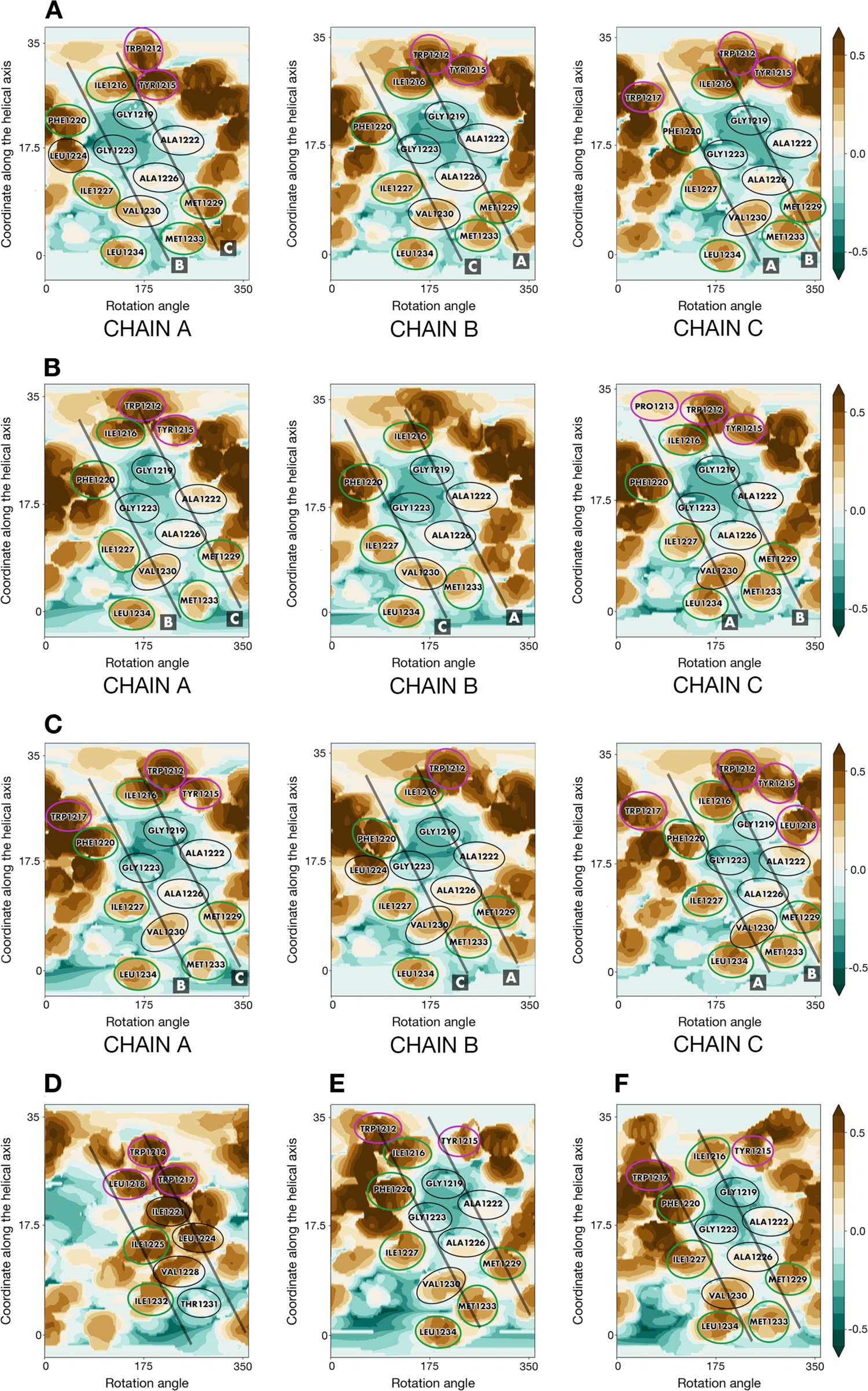
Helix/helix interfaces in *S_OPT* and certain alternative models on MHP maps. (A) *S_OPT* after energy minimisation. (B) *S_OPT* at 1μs in trajectory run1. (C) *S_OPT* at 1μs in trajectory run2. (D-F) Models created by Casalino et al. (2020) (D) and two versions (E and F) of the S-TMD by Woo et al. (2020) built using HIV’s gp41 as template. Identical positions are encircled in purple, conservative and semi-conservative residues are encircled in green, non-conservative residues present on the helix/helix interface are encircled in black. In the MHP map for each chain, schematic projections of the other two chains are shown as thick semi-transparent black lines and, where appropriate, are designated by letters A to C. For other details see legend to Fig. S1.

**Figure S4.**
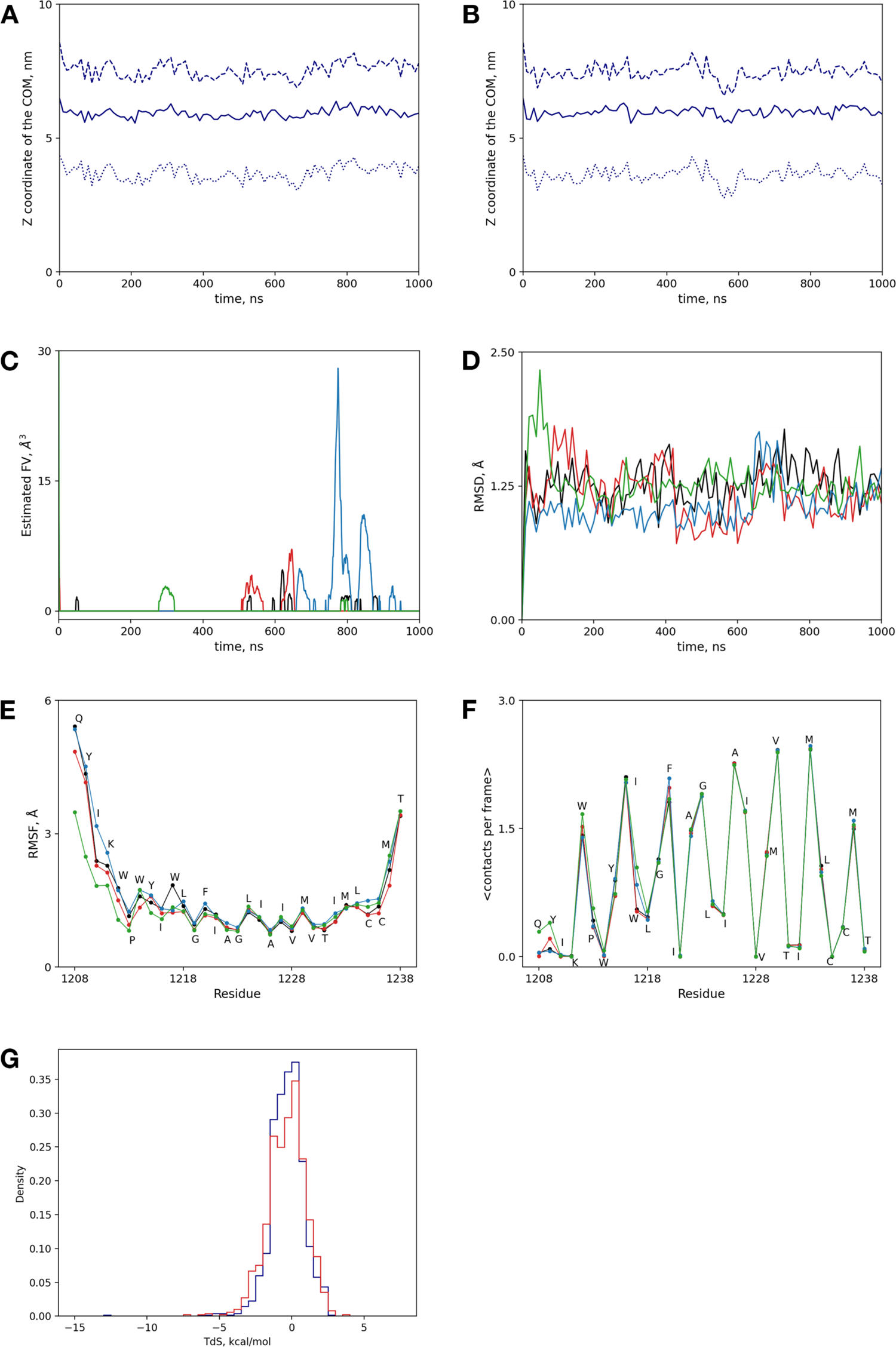
Performance of *S_OPT* and *S_OPT-PLM* in a POPC bilayer. (A-B) Z coordinate of the centre of mass (COM) of *S_OPT* residues 1212-1234 in run1 (A) and run2 (B), indicating trimer localization (solid lines) in the membrane with respect to the COM of the P atoms of the upper (dashed lines) and lower (dotted lines) leaflets of the POPC bilayer. (C) RMSD calculated for the backbone atoms of residues 1212-1234 in *S_OPT* (black and red) and *S_OPT-PLM* (blue and green). Frame 0 was used as the reference structure in all cases. (D) Estimated free volume in the lumen in *S_OPT* (black and red) and *S_OPT-PLM* (blue and green) trimers over the course of their respective trajectories. (E) RMSF values for individual residues in *S_OPT* (black and red) and *S_OPT-PLM* (blue and green). (F) Involvement of individual residues in helix/helix non-covalent interactions in palmitoylated (blue and green) and unpalmitoylated (black and red) variants of *S_OPT*. (G) Normalised histograms of TΔS distribution for POPC molecules in systems containing *S_OPT* (blue) and *S_OPT-PLM* (red) as compared to a “pure” POPC bilayer.

**Figure S5.**
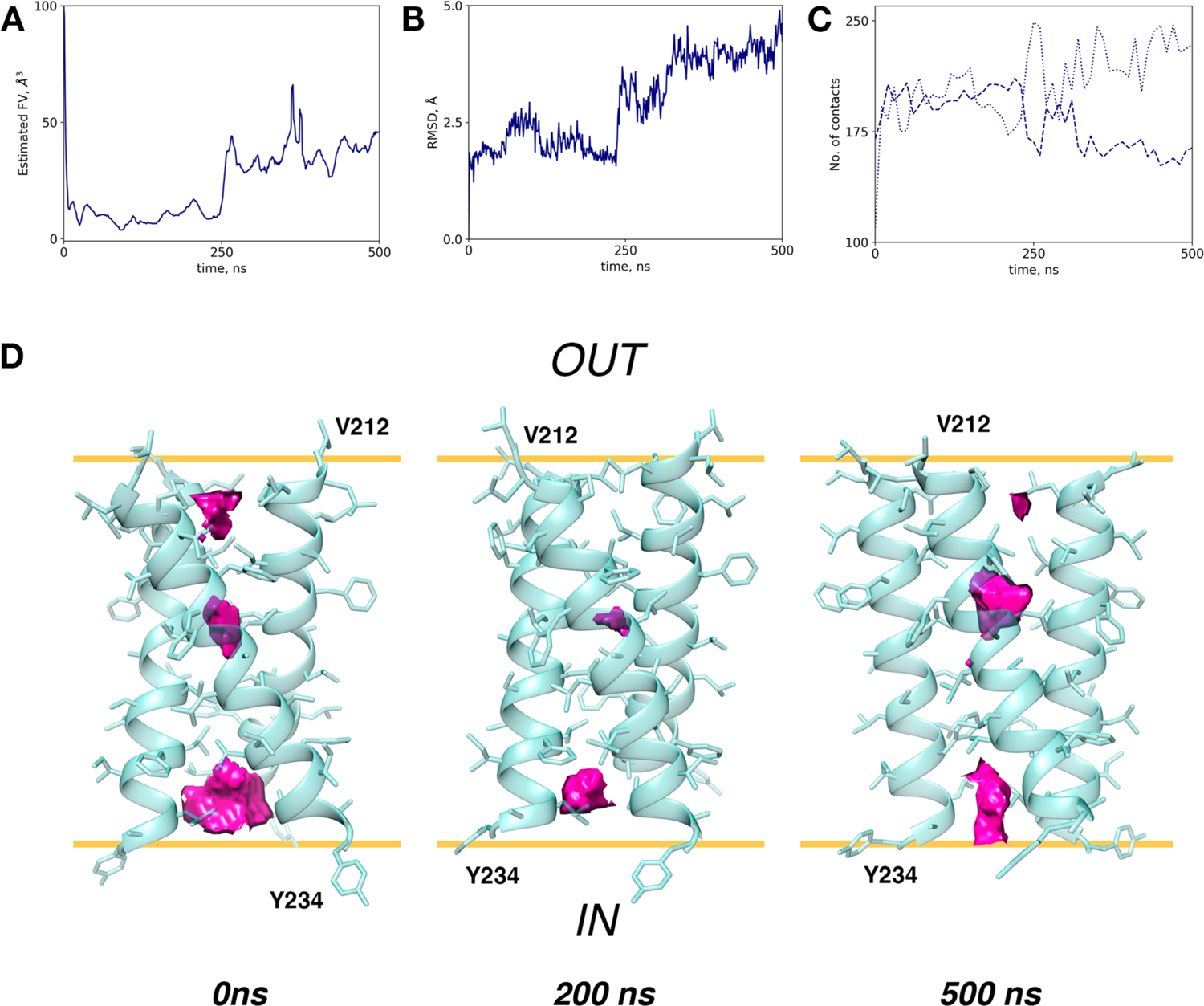
Results of the MD simulation of TNFR-1 TMD in a POPC bilayer. (A) Estimated free volume inside TMD during an MD simulation in a POPC bilayer. (B) RMSD for the backbone atoms (residues 212 to 233). (C) Protein/protein (dashed line) and protein/lipid (dotted line) contacts for the side chains constituting the interface (214, 217, 218, 219, 221, 222, 224, 225, 228, 229, 232). (D) Visualisation of estimated free volume (pink) inside the TMD at 0ns, 200ns and 500ns (averaging over 200 frames not applied). 3D models are shown in cartoon/stick representation. Approximations of carbonyl/ester oxygen planes are shown as yellow lines, while the inside of the cell and the extracellular space are labelled “IN” and “OUT”, respectively.

**Figure S6.**
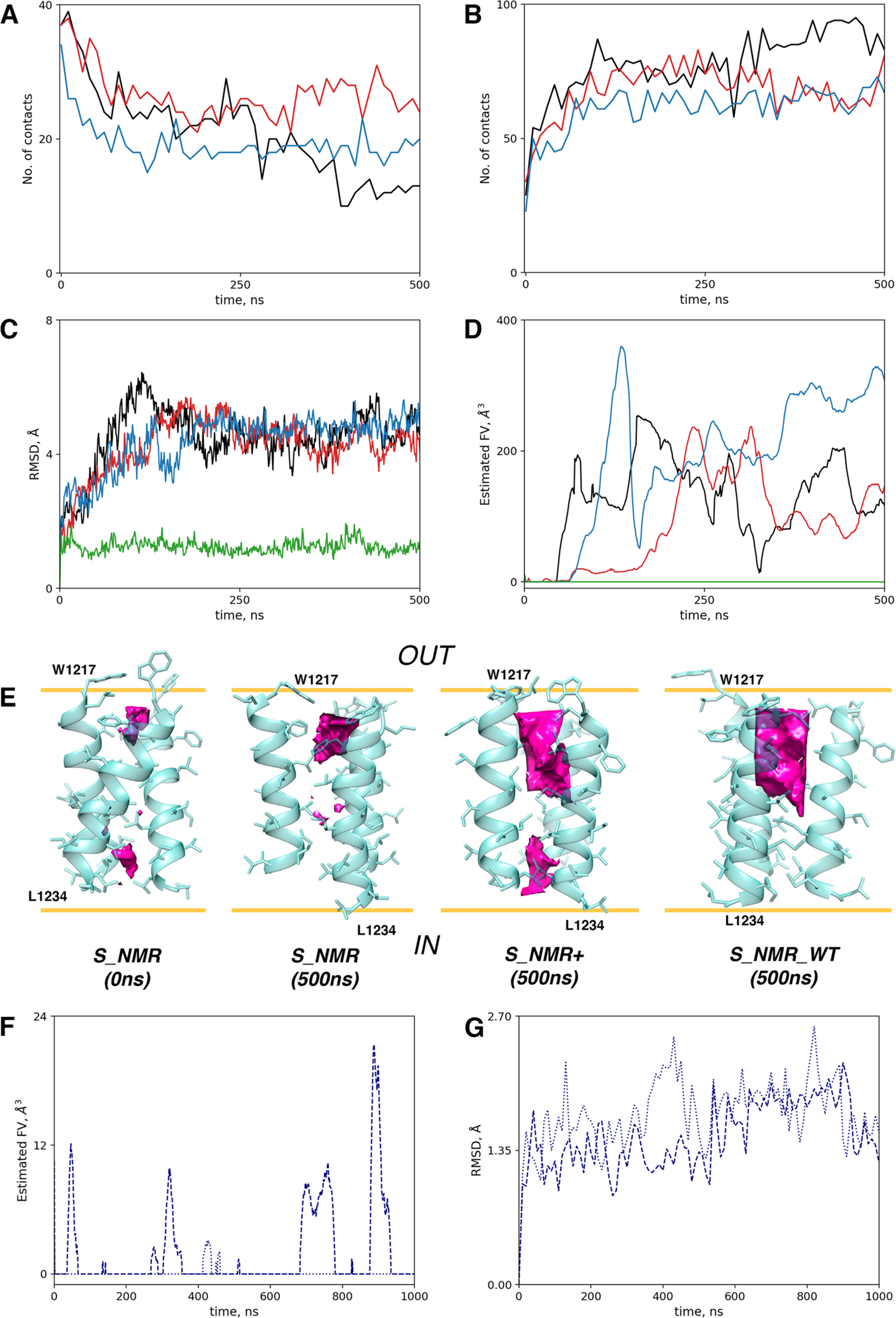
Performance of NMR-based models and mutant forms in a POPC bilayer. (A-B) Contacts made by the side chains of I1221, I1225, L/M1229 and L/M1233 with lipids (A) and with the same side chains in other helices of the homotrimer (B) calculated for trajectories *S_NMR* (black), *S_NMR+* (red) and *S_NMR-WT* (blue). (C-D) RMSD of the backbone atoms of the proposed TMD (residues 1218-1234) and estimated free volume inside the lumens model trimers *S_NMR* (black), *S_NMR+* (red), *S_NMR-WT* (blue) and *S_OPT* (green) over the course of MD simulations. (E) Visualisation of FV in model trimers *S_NMR*, *S_NMR+* and *S_NMR-WT*. 3D models are shown in cartoon/stick representation, and free volume is rendered as pink slabs (averaging over 200 frames not applied). Approximations of carbonyl/ester oxygen planes are rendered as yellow lines, the inside of the virion and the external environment are labelled “IN” and “OUT”, respectively. (F) Estimated free volume in the lumen of *S_OPT-LC* (dashed line) and *S_OPT-LS* (dotted line) over the course of their respective trajectories. (G) RMSD calculated for the backbone atoms of residues 1212-1234 in model trimers *S_OPT-LC* (dashed line) and *S_OPT-LS* (dotted line); frame 0 was used as the reference structure in both cases.

**Figure S7.**
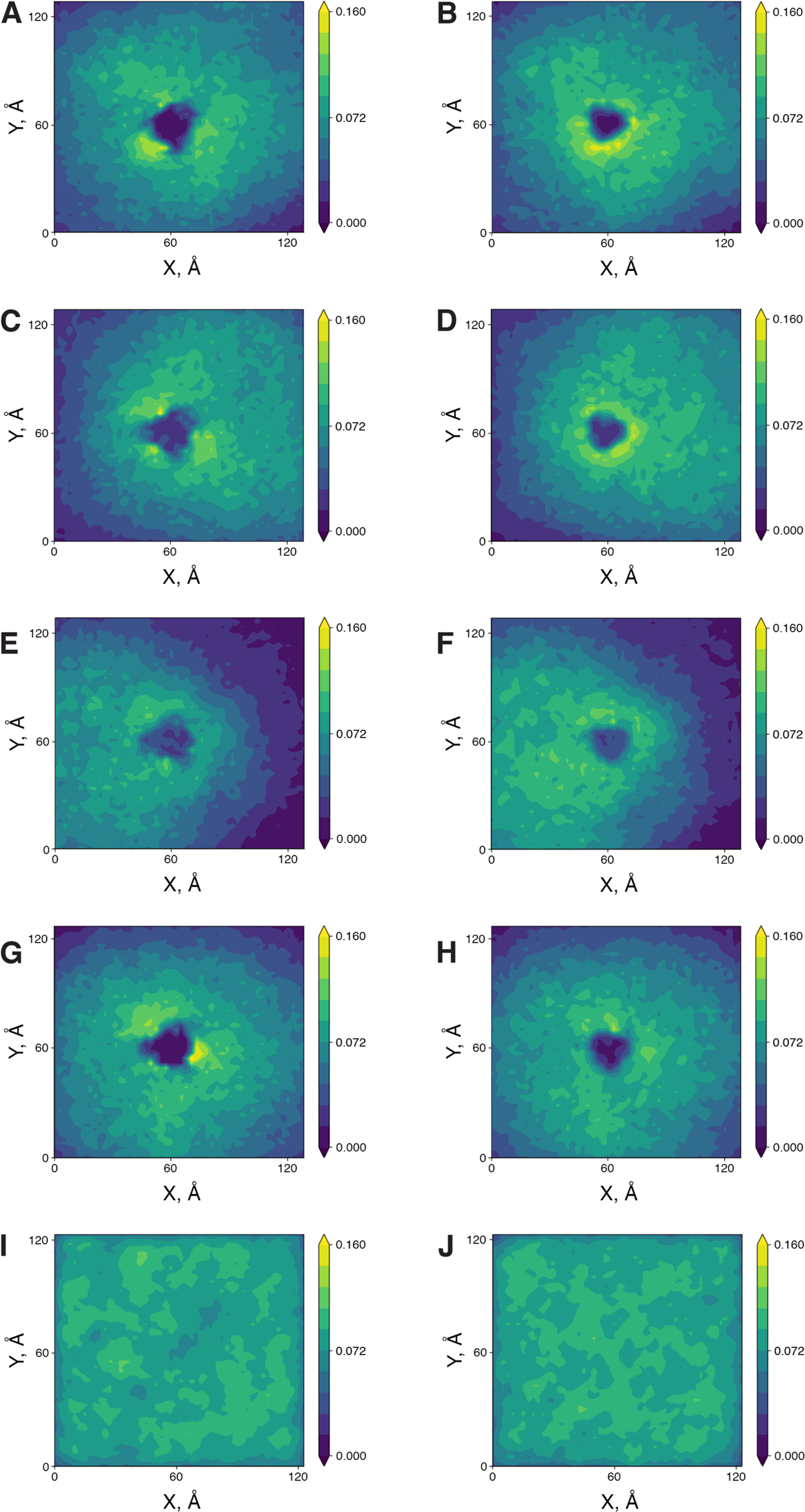
Order parametre maps plotted for model POPC bilayers. (A-B) *S_OPT* run 1 top leaflet (A) and bottom leaflet (B). (C-D) *S_OPT* run 2 top leaflet (C) and bottom leaflet (D). (E-F) *S_OPT-PLM* run 1 top leaflet (E) and bottom leaflet (F). (G-H) *S_OPT* run 2 top leaflet (G) and bottom leaflet (H). (I-J) A “pure” POPC bilayer top leaflet (I) and bottom leaflet (J). Maps were plotted over a period from 50ns to 1000ns for each trajectory. All bilayers are oriented in such a way that the normal to the bilayer plane is parallel to the Z axis, and the axes in maps, labelled X and Y, are oriented along eponymous axes in each system. An S_CD_ order parameter colour scale is presented on the right of each map.

**Figure S8.**
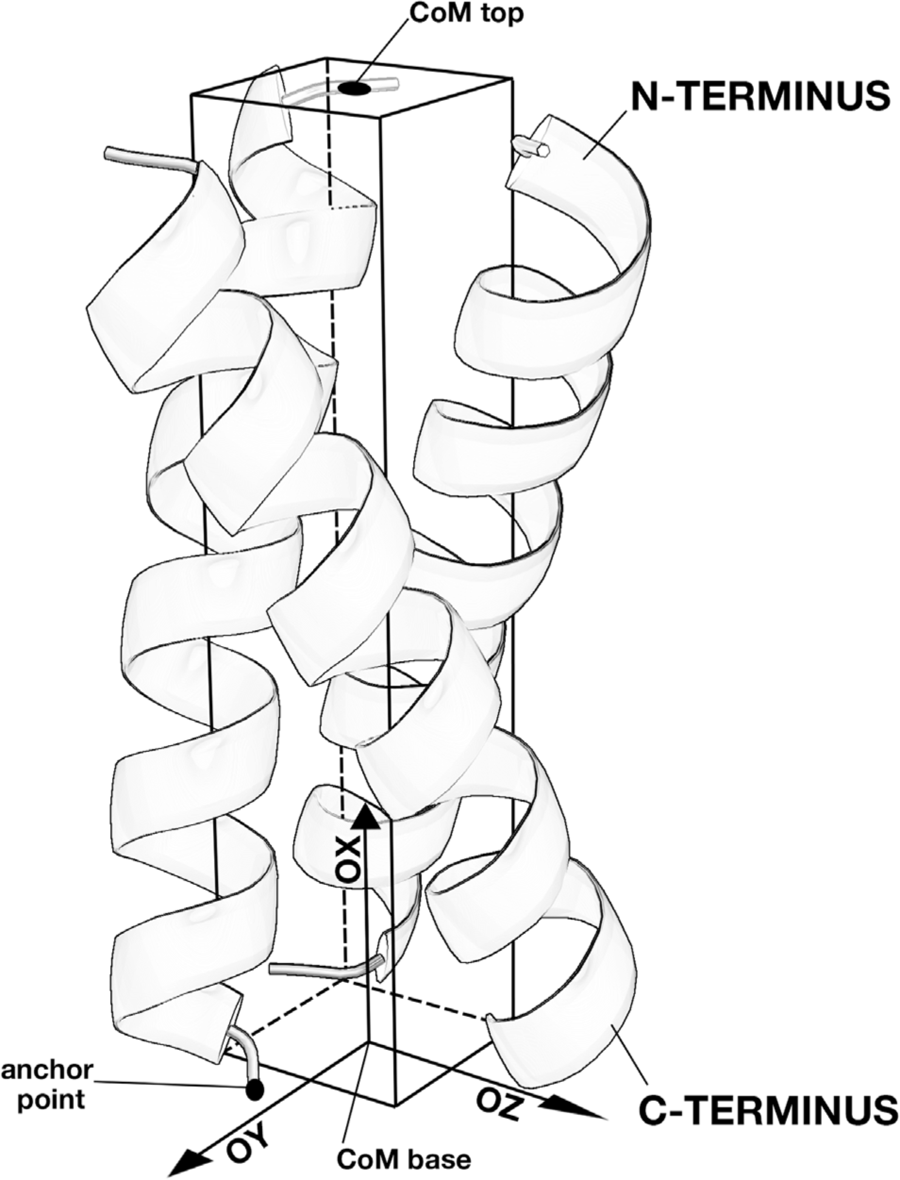
Mesh applied to calculate free volume (FV) inside a helical trimer. The helices are shown in ribbon representation. The confines of the mesh are shown as black lines. The CoM top and anchor point are shown as black dots. Axes of the local coordinate system originating at CoM base are shown conventionally and labelled accordingly. See “Methods” for details.

